# Microglia-Specific Molecular Magnetic Resonance Imaging Probe Enables Noninvasive Separation of Parkinsonian Mice from Controls

**DOI:** 10.64898/2026.06.23.734093

**Authors:** Xianwei Sun, Andrew Badachhape, Terry-Elinor Reid, Esther Ngan, Sheelu Monga, Jeannie Chin, Ananth Annapragada, Henry Lowe, Ngeh Toyang, Eric Tanifum

## Abstract

Neuroinflammation mediated by reactive microgliosis is a central driver of Parkinson’s disease (PD) pathogenesis. This inflammatory process unfolds years before clinical symptoms, creating an opportunity for early intervention. *In vivo* imaging technologies that could detect and quantify microglial reactivity are therefore essential for early diagnosis, patient stratification, and evaluating emerging immunomodulatory therapies that target this fundamental driver of PD progression. Yet no standardized, sensitive, and specific technology currently achieves this goal. Molecular magnetic resonance imaging (mMRI) is uniquely suitable to address this problem because it integrates inherent high spatial resolution and soft-tissue contrast of conventional MRI with molecularly targeted contrast agents, enabling simultaneous acquisition of anatomical detail and functional/biological information at submillimeter isotropic resolution. Here we present a novel mMRI probe designed to specifically target colony stimulating factor-1 receptor, expressed primarily on microglia in the brain. *In silico* data show that the targeting ligand binds the extracellular Ig domain of the receptor. In *vitro* cell uptake studies with both murine and human microglia cell lines show that the probe binds the receptor triggering active cell uptake and *in vivo* MRI enabled effective separation of the A53T mouse model of Parkinson’s disease from control mice using radiomics-assisted MR image analysis. *Ex-vivo* immunohistochemical analysis showed signal from the probe largely in the cytosolic compartment of IBA-1 reactive cells, confirming that the observed *in vivo* MRI signal is due primarily to retention of the agent by microglia. This novel technology has the potential to interrogate the rgional presentation of microglial activation in PD.

**One Sentence Summary:** A microglia targeted MRI probe generates disease-specific contrast after injection, clearly distinguishing A53T Parkinsonian mice from controls.

## INTRODUCTION

Parkinson’s disease (PD) is the fastest growing neurological disorder, affecting over 11.8 million individuals across the globe in 2021 (1). In the United States, about 90,000 new cases are reported annually, representing a 50% increase from previous estimates, leading to an estimated 1.2 million people living with the disease by 2030 (2). PD is also highly heterogeneous (3) with complex pathogenesis (4) that results in significant neuronal damage occurring before motor deficits appear (5). Because current diagnostic metrics focus on motor symptoms, clinical diagnosis is delayed relative to underlying pathophysiology. This complexity also limits the development of disease modifying treatments as no single biomarker captures the disease’s multifaceted biology and therapies aimed at one mechanism cannot overcome a disorder driven by many interacting pathways. Technologies that enable earlier detection and biomarker-guided stratification are required to accelerate the development of therapies that can truly alter the disease course.

Neuroinflammation mediated by microglia activation is increasingly recognized as a central and dynamic contributor to the pathogenesis of PD, making it a potential biomarker candidate (6–8). In prodromal and early stages of the disease, microglia may initially adopt a protective phenotype, clearing α-synuclein aggregates and releasing trophic factors. However, as the disease progresses, microglia shift toward a chronic pro-inflammatory state accompanied by reactive astrocytosis (9–12). Persistent glial activation accelerates neuronal death as accumulation of α-syn aggregates in the form of Lewy bodies and Lewy neurites spreads from the medulla oblongata and olfactory systems through to the lower raphe nuclei and locus, to the substantia nigra pars compacta and amygdala, where it triggers the first clinical symptoms (13). Furthermore, persistent glial activation perpetuates recruitment of peripheral immune cells into the brain where they amplify neuroinflammation and contribute to disease progression by altering T-cell behavior, monocyte trafficking, and peripheral cytokine profiles (14). Taken together, these findings highlight activated microglia as early sensitive biomarkers of disease because they are early dynamic responders to α-synuclein pathology (15), act as central amplifiers of neuroinflammation through cytokine signaling (16–18), and bridge innate and adaptive immunity (19). This suggests that noninvasive imaging tools that can profile microglia activity *in vivo* could enable earlier detection and patient stratification for trials.

Positron emission tomography (PET) tracers, the current gold standard molecular imaging tools used to noninvasively profile amyloid-β plaques and hyperphosphorylated tau tangles, have been immensely useful in Alzheimer’s disease. However, such tools have not been adopted for routine clinical diagnosis or monitoring of neuroinflammation or Lewy pathology in PD. PET tracers of the mitochondrial translocator protein (TSPO) have provided *in vivo* evidence of microglial activation in PD patients (*20, 21*), but variability in tracer specificity and genetic polymorphisms affecting TSPO binding (22) have limited their use to research protocols. Alternative approaches including second- and third-generation TSPO tracers with improved signal-to-noise ratios (20) and exploratory PET ligands targeting alternative neuroinflammatory pathways including COX-2, P2X7 receptors, and the membrane bound protein, colony-stimulating factor 1 receptor (CSF1-R) (*23, 24*), are being actively pursued to overcome limitations of TSPO. Tracers targeting CSF1-R such as [¹¹C]CPPC and [¹⁸F]CPPC have shown promising preclinical and early clinical results, but challenges remain around optimizing brain penetration, selectivity, and low signal-to-noise (25–27) for accurate noninvasive spatiotemporal routine profiling of microglia-mediated neuroinflammation in the clinic. Previously, in rodent models of neurodegenerative disorders, we demonstrated that following intravenous administration (i.v.), liposomal nanoparticles bearing a magnetic resonance imaging (MRI) contrast payload and labeled with specific ligands passively cross into the brain parenchyma and label specific disease biomarkers based on the targeting ligand. These tools enabled *in vivo* separation of treated disease cohorts from controls using noninvasive molecular magnetic resonance imaging (mMRI) (*28–31*).

Based on our recent discovery that the flavonoid Caflanone is a potent inhibitor of CSF1-R (*32*), we hypothesized that a Caflanone-labeled variant of our mMRI probe, NanoCaflanone diagnostic (NCDx), would enable selective microglial binding and trigger cell-specific receptor-mediated internalization of the probe. CSF1-R’s high enrichment on CNS-resident myeloid cells and its dynamic upregulation during neuroinflammation (*33*) could provide both specificity and disease-responsive uptake. Combined with the high-resolution of MRI, the probe could highlight brain regions where deposition of Lewy pathology is accompanied by microglial activation, enabling noninvasive separation of disease cohorts from controls. In this study, we demonstrate that a polyethylene glycol (PEG)-tethered analogue of Caflanone (PEGylated Caflanone) binds the extracellular domain of CSF1-R and that NCDx actively labels murine and human microglia cell lines *in vitro*. We further demonstrate that following i.v. administration in the A53T α-syn transgenic line M83 mouse model of PD, NCDx also labels microglia *in vivo*, enabling noninvasive separation of pathology-bearing transgenic mice from wildtype controls. *Ex-vivo* immunohistochemistry (IHC) validates these *in vivo* data. Our findings demonstrate that NCDx targets microglia, enabling noninvasive separation of Parkinsonian mice from controls *in vivo*, and has the potential to profile microglia-mediated neuroinflammation in other central nervous system disorders.

## RESULTS

### A PEGylated analogue of Caflanone binds the extracellular domain of CSF1-R

The targeting ligand on our previously reported mMRI probes (*31*) is expressed on a PEG chain, therefore we started by establishing that a PEGylated analogue of Caflanone (Fig. **1A-B**) has a suitable binding site on the extracellular domain of CSF1-R (Fig. **1C**) using molecular simulation studies. We prioritized sites on the extracellular domain because our nanoparticle requires specific surface interactions with the target cells for optimal efficacy. Specifically, we computationally explored possible druggable binding pockets on the extracellular immunoglobin (Ig)-like domain of CSF1-R using the Schrödinger tool, SiteMap (34). This resulted in four potential binding sites (Fig. **1D**), including Site 1 (SiteScore of 0.920), Site 2 (SiteScore of 0.912), Site 3 (SiteScore of 0.916), and Site 4 (SiteScore of 0.913). Using molecular simulation docking studies, two sites (2 and 3) emerged as potential high affinity binding sites to the molecule. Site 3, located on Ig D5 had the lowest predicted binding energy of -8.569 kcal/mol, indicating high affinity selective binding to the site (Fig. **1E**). In this scenario, the construction extended the surface of the Ig D5 region with the amphiphilic PEG group traversing a narrow hydrophobic tunnel in which the nitrogen of the carbamate linker made H-bond interactions with the Cys 485 backbone and another H-bond between an oxygen atom in the PEG unit and Trp 496 residue. The flavonoid core structure on the other end showed H-bonds and a salt-bridge interaction with Lys 465 residue, extending the conformation of the construct and further anchoring it to the site. Site 2 (Fig. **1F**), located between the Ig D2-D3 interface and close to the binding site of the natural ligand (CSF1), had the 2^nd^ best glide score of -7.132 kcal/mol. In this scenario, the PEG group interacted with a more solvent-exposed region of Ig D2, forming H-bonds with the residues of Arg 166 and Lys 168, resulting in a 109° angular bend of PEG at C-1 of the carbamate. The flavonoid core extended into a hydrophobic cavity, making H-bonds with the backbone of Arg 66 and Phe 67, and multiple H-bonds with Gln 81.

**Fig. 1.**
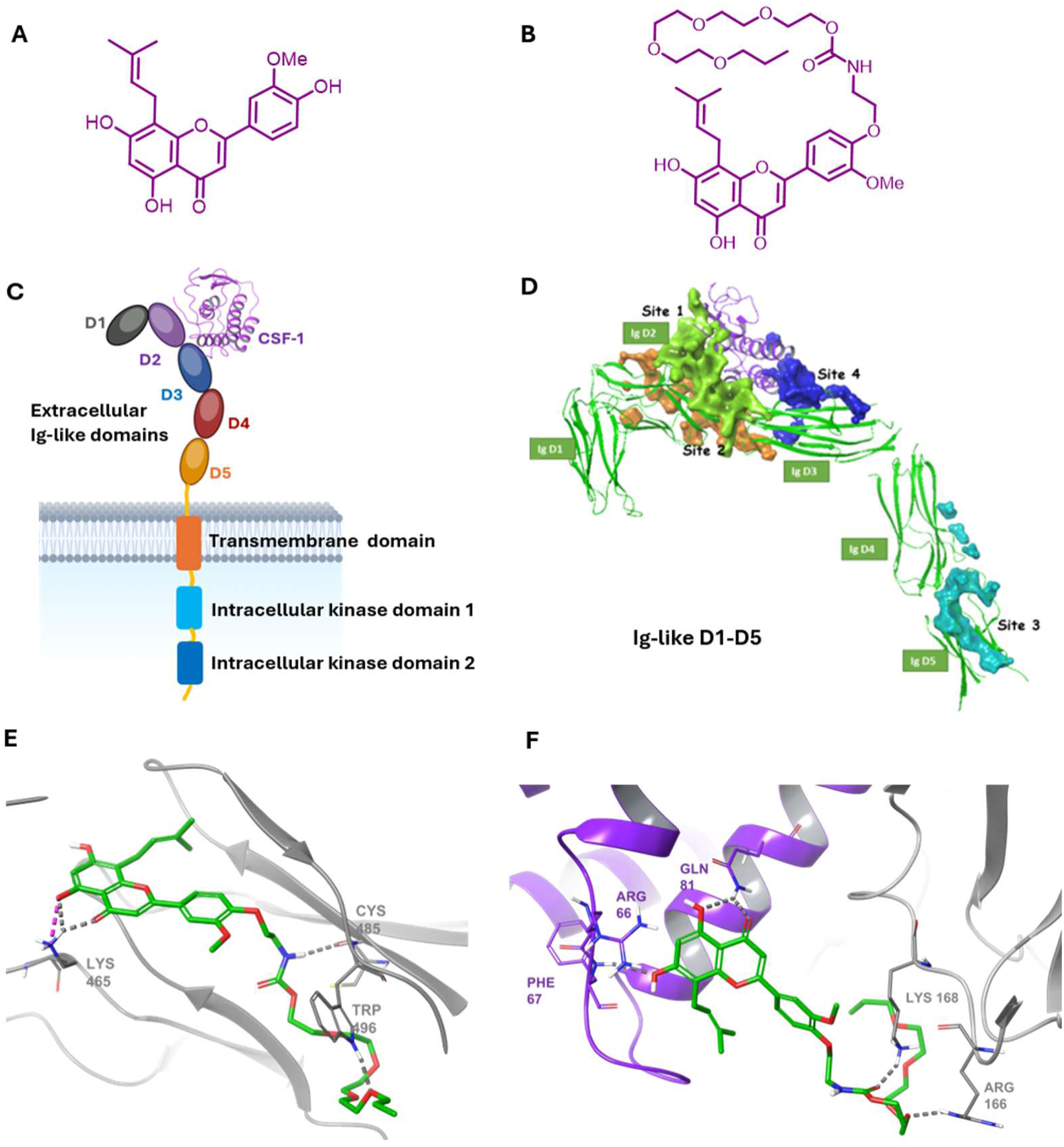
PEGylated Caflanone interacts with the extracellular Ig domains of CSF1-R. The molecular structures of (**A**) Caflanone and (**B**) PEGylated Caflanone. (**C**) Schematic of the CSF1-R transmembrane receptor illustrating the intracellular kinase domain, the transmembrane domains, and the 5 extracellular Ig-like domains (D1-D5) with bound CSF1; **D**) Four druggable sites on the extracellular Ig-like domains with site 1 (green) and site 2 (orange) on D2, site 3 (cyan) on D5, and site 4 (blue) on D3; Predicted interaction of PEGylated Caflanone with CSF1-R when docked (**E**) in site 3 or (**F**) in site 2.

### CSF1-R targeted mMRI probe (NCDx) designed with 4 key components

Based on our *in silico* data showing that PEGylated Caflanone binds the extracellular Ig-like domain of CSF1-R, we designed and formulated a Caflanone labeled mMRI probe (NCDx). Key chemical components (Fig. **2**) included: (a) a novel Caflanone-PEG(3400)-DSPE conjugate as the targeting moiety, prepared as described in the supporting materials section; (b) Gd(III)DOTA-DSPE as the MRI contrast source synthesized using a previously reported synthetic protocol (30), (c) mPEG(2000)-DSPE obtained from commercial sources to impart long systemic circulation half-life; and (d) Commercial Dil dye as a fluorescent reporter on the probe. Following assembly of all reagents, we formulated the nanoparticles using classical liposome hydration and extrusion protocols (30) to obtain a formulation with a particle hydrodynamic diameter of 148.95 ± 26.63 nm using Dynamic Light Scattering as shown in the supporting materials section (fig. **S1**) and a total lipid molarity 52.82 ± 3.68 mM, Gd(III) concentration of 17.16 ± 1.16 mM using Inductively Coupled Plasma mass spectrometry and a systemic circulation half-life of 11.83 h (R^2^ = 0.91, Sy.x = 0.03) (fig. **S2**).

**Fig. 2.**
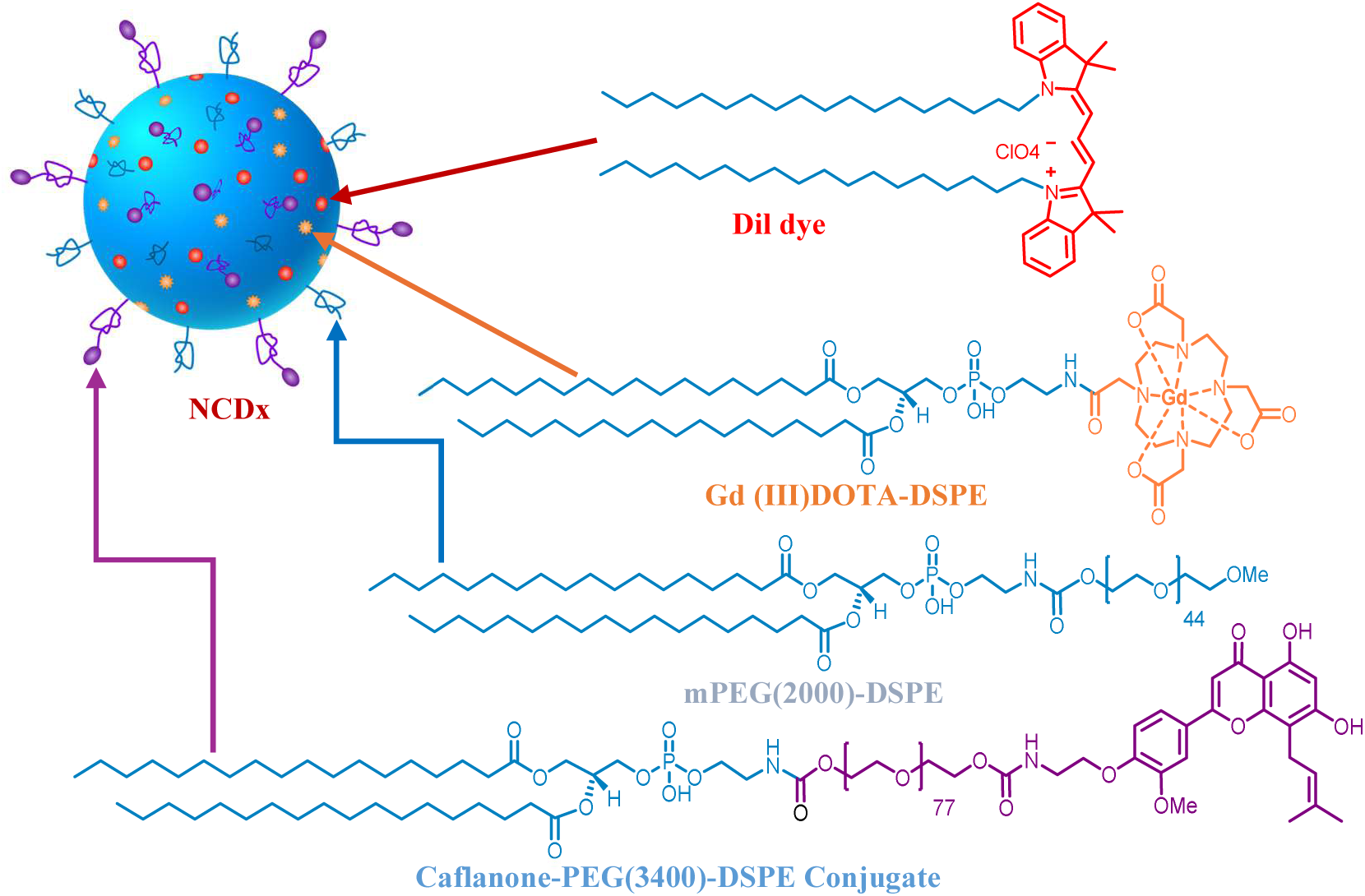
Key molecular components of Caflanone-labeled mMRI probe, (NCDx).

### CSF1-R expressing cells actively internalize NCDx upon exposure to the probe

To verify that NCDx specifically interacts with CSF1-R expressing cells and triggers internalization, we performed *in vitro* cell uptake experiments using murine (SIM-A9) and human (HMC-3) microglia cell lines. Briefly, cells were incubated with NCDx for 2 hours followed by a wash off to remove any uninternalized particles. Controls included cells treated with a nontargeted variant of the probe (NT) and untreated cells (UT). Using confocal microscopy of mounted cells (Fig. **3A**), we found that treated (NCDx) SIM-A9 cells showed strong NCDx signal. Cytoskeletal imaging using Phalloidin, confirmed NCDx signal within the cytosol. 3D visualization of the image accentuated cell borders, unequivocally demonstrating that the NCDx signal was cytosolic. Importantly, internalization was seen within two hours of exposure. Neither NT nor UT controls showed evidence of NCDx internalization. We next assessed the effect of CSF1 cell activation on NCDx uptake. Indeed, CSF1 activation resulted in a statistically significant increase in NCDx uptake by SIMA-9 cells (CSF1 + NCDx) compared to nonactivated cells (NCDx) after 2 hours (Fig. **3B**). To determine the time course of uptake, we assessed change in pixel intensity per cell every 15 minutes over 4 hours (Fig. **3C**), and it showed saturation after 90 minutes of incubation. We observed similar results using HMC-3 cells and Raw 264.7 cells from cell uptake studies (Fig. **3D** and fig. **S3A**, respectively), cell activation with CSF1 (Fig. **3E** and fig. **S3B,** respectively) and time dependent uptake (Fig. **3F** and fig. **S3C,** respectively). As expected, because they do not express CSF1-R, murine and human primary astrocytes subjected to the same treatment protocols did not show evidence of internalization when exposed to NCDx under similar conditions. These data suggest that the targeting ligand (Caflanone) on NCDx plays a role in the internalization of the nanoparticles.

**Fig. 3.**
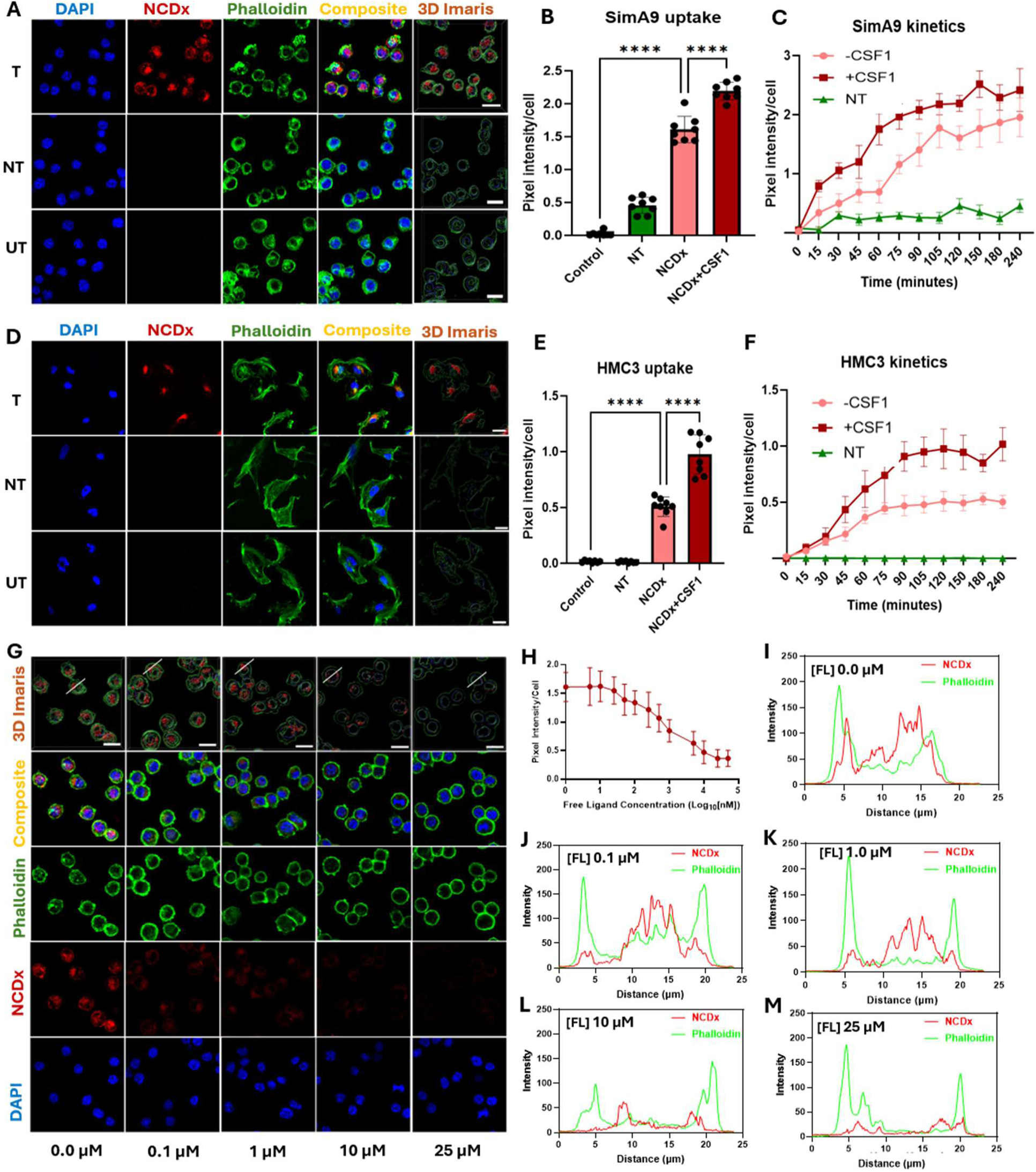
Murine and human microglia cell lines actively internalize NCDx. (**A**) SIM-A9 cells (murine microglia cell line) show avid internalization of NCDx within 2 hours of exposure; (**B**) Activation of SIM-A9 cells with CSF1 results in increased uptake of NCDx; (**C**) A plot of signal intensity per cell against exposure time demonstrating that **NCDx** internalization exhibits saturation kinetics; (**D**) HMC-3 cells (human microglia cell line) also show avid uptake of **NCDx**; (**E**) Activation of HMC-3 with CSF1 results in increased uptake of NCDx; (**F**) NCDx uptake by HMC-3 cells exhibit saturation kinetics; **(G**) Confocal microscopy images show that blocking the receptor binding sites with the free ligand inhibits NCDx uptake; (**H**) A plot of NCDx signal intensity per cell co-incubated with the free ligand showing NCDx signal decay with increasing concentration of the free ligand. (**I** to **M**) Intensity curve along the indicative profile line in confocal images. Scale bar = 20 µm.sss

Finally, to establish particle uptake specificity, we performed a blocking study in which cells were incubated with NCDx at increasing concentrations of free ligand. Increasing free ligand concentration correlated with diminished NCDx uptake (Fig. **3G**, **H** and fig. **S4**). To further quantify free ligand blocking efficiency, we conducted an ImageJ RGB profiler analysis of NCDx fluorescence intensity at each concentration (Fig. **I-M** and fig. **S5**). We observed a progressive reduction in NCDx fluorescence signal (red) as free ligand concentration increased. Taken together, these in *vitro* cell uptake data suggest that CSF1-R expressing cells actively internalize NCDx.

### Conventional *in vivo* mMRI showed a significant difference in percent signal change distribution between TG and WT

Based on our *in vitro* data demonstrating rapid, specific internalization, we next interrogated the potential of NCDx to noninvasively separate Parkinsonian brains from controls. Towards this, we leveraged the transgenic A53T α-synuclein (α-syn) M83 PD mouse model for which Lewy body pathology development parallels disease progression in humans; changes appear first in the brainstem and olfactory systems then the midbrain and eventually cortical areas^35^. Eight 12 - 14-month-old mice transgenic mice (TG) underwent pre-contrast (PRE) scans using an axial T1-weighted spin echo (T1w-SE) protocol to obtain baseline images, immediately followed by injection of NCDx (dose, 0.15 mmol Gd(III)/kg). 4 days later, when unbound contrast had cleared systemic circulation, the mice were subjected to the same scan protocol to obtain post-contrast (POST) images. Pseudo colored coronal sections from cortices and olfactory bulbs of transgenic mice (Fig. **4A**) showed an increase in signal from the PRE to POST images. The same observation was not apparent in wild type mice (WT) controls injected with NCDx (Fig. **4B**). Average change in signal intensity between PRE and POST images, calculated by subtracting the signal intensity in the PRE image from the POST image for corresponding brain sections, showed a significantly larger increase for the TG mice compared to WT in both brain regions (Fig. **4C****, 4D**). A panel of all cortices and olfactory bulbs of all treated mice and controls are shown in fig. **S6** in the supporting materials.

**Fig. 4:**
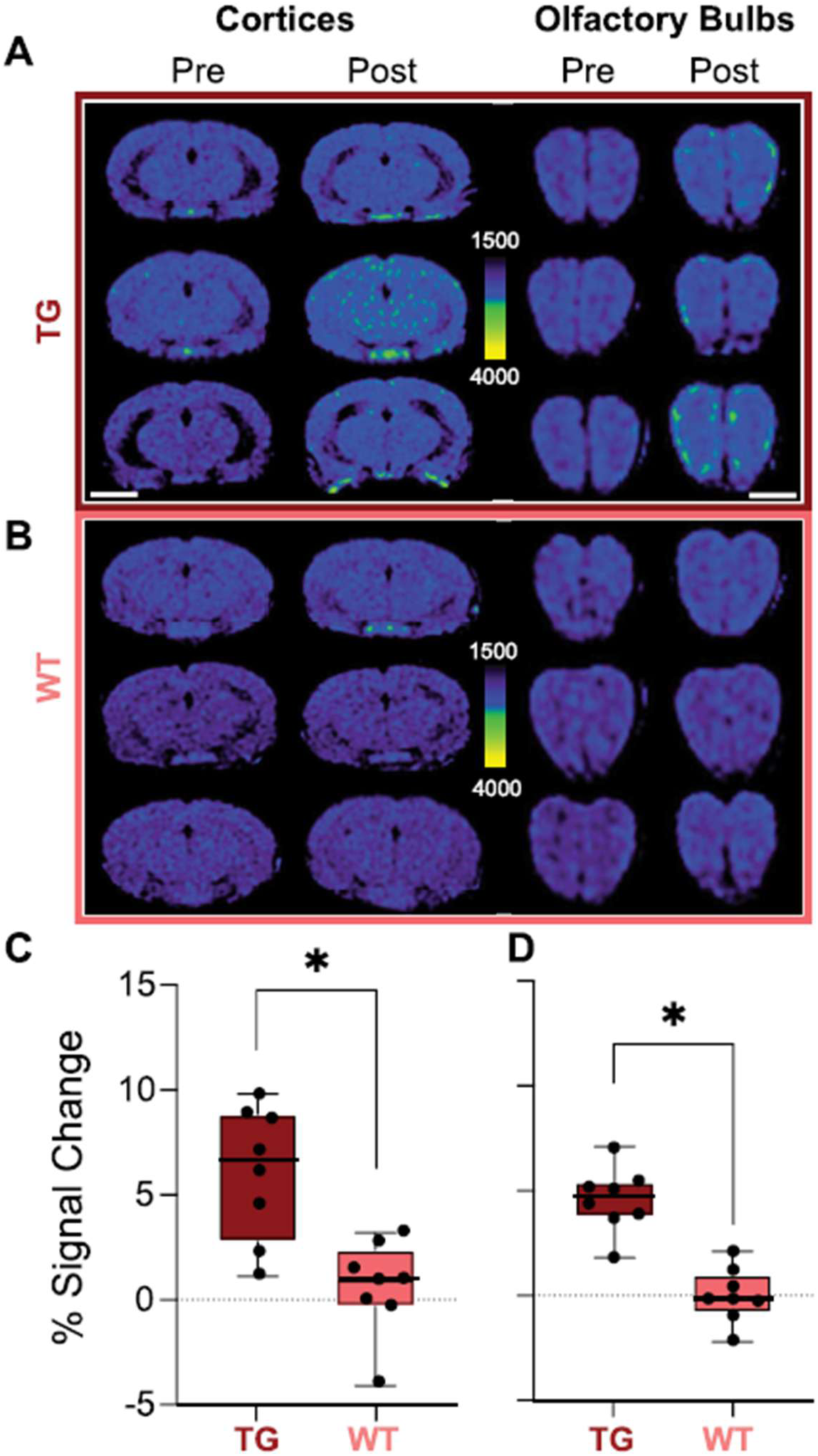
In *vivo* contrast-enhanced MRI of mice injected with NCDx. (**A**) Axial 3.6 mm thick slabs cortices (Bregma 0 mm) and olfactory bulbs (Bregma 4-5 mm) of TG mice using a jet colormap to illustrate particle label intensity show apparent increase in signal intensity in POST scans compared to PRE. (**B**) Similar sections from WT mice subjected to the same experimental protocol do not show any apparent signal change in PRE versus POST scans. All images were analyzed on the same MR signal intensity scale (1500-4000 a.u.); (**C**) Box and whisker plot quantification of percent change in signal intensity demonstrate statistically signal increase in TG versus WT mice cortices; (**D**) Box and whisker plot quantification of percent change in signal intensity demonstrate statistically signal increase in TG versus WT mice olfactory bulbs. Wilcoxon rank sum test: *p<0.05. Scale bar = 3 mm.

### Machine learning (ML) enabled full separation of transgenic mice from wild types

Although percent change in signal intensity statistically differentiated TG from WT mice, not all TG mice showed a greater change than the WT mice in both cortices and olfactory bulbs. Therefore, we employed machine learning (ML)-assisted image analysis to determine if subtle, micro-scale probe-related changes in the *in vivo* post-contrast MRI data could fully distinguish TG from WT mice. Specifically, we incorporated radiomic features (RF) that can quantify MRI data by capturing intensity distributions, spatial textures, geometric properties, and higher-order signal transformations that reflect underlying tissue biology over the entire regional volume with far greater granularity than conventional ROI signal intensity analysis (*35, 36*).

A total of 1284 radiomic features (RFs), including both filtered and unfiltered, for each brain per region (cerebral cortex or olfactory bulb) were extracted. Following feature selection, the most informative RFs from each brain region were retained for classification. First, we ran the analysis using raw image data because this is the most straight forward. A multi-layer perceptron (MLP) classifier with only three cortex RFs was sufficient to achieve perfect performance, with validation and testing accuracy and ROC values reaching 1.0. Using olfactory bulb RFs allowed the model complexity to be reduced to logistic regression, albeit with a total of 10 RFs (Table **1**). To minimize potential bias in RF computation associated with voxel anisotropy and to conform to established radiomics recommendations (^38^), we also ran our analyses using isotropically resampled images. In these scenarios, a MLP classifier using three cortex RFs and a random forest classifier with 3 olfactory bulb RFs achieved perfect classification using fewer features than with raw data. Finally, to remove instrument artifacts, we pixel-normalized the isotropically resampled images and re-ran the analyses. Perfect classification was now achieved using a logistic regression classifier with a single cortex RF. However, performance decreased when considering cortex and olfactory bulb together. Overall, images resampled to isotropic resolution achieved perfect classification to separate TG mice with Parkinson’s disease pathology from WT controls using the fewest features and simplest classifiers. Further, classifiers using olfactory bulb RFs used simpler models but required a larger number of features (Table **1**). For all analyses, when RFs from both the cortex and olfactory bulb were combined, the models required more features and/or exhibited poorer performance.

**Table 1.** Classification of in vivo MR images of transgenic mice from wild types using Radiomic features. LDA = linear discriminant analysis; KNN = k-nearest neighbor; MLP = multilayer perceptron neural network; AUC = Area under the Rceicer Operating Characteristic Curve.

| Image type | ROI(s) | Best classifier | Validation Accuracy | Validation AUC | Testing Accuracy | Testing AUC | Number of selected RFs |
| --- | --- | --- | --- | --- | --- | --- | --- |
| raw | cortex | MLP | 1 | 1 | 1 | 1 | 3 |
| raw | olfactory bulb | logistic | 1 | 1 | 1 | 1 | 10 |
| raw | cortex and olfactory bulb | MLP | 1 | 1 | 1 | 1 | 5 (2 from cortex) |
| isotropic | cortex | MLP | 1 | 1 | 1 | 1 | 2 |
| isotropic | olfactory bulb | Random forest | 1 | 1 | 1 | 1 | 3 |
| isotropic | cortex and olfactory bulb | Gaussian naive bayes | 1 | 1 | 1 | 1 | 11 (5 from cortex) |
| isotropic and normalized | cortex | logistic | 1 | 1 | 1 | 1 | 1 |
| isotropic and normalized | olfactory bulb | LDA | 1 | 1 | 1 | 1 | 4 |
| isotropic and normalized | cortex and olfactory bulb | KNN with k=1 | 1 | 1 | 0.75 | 0.75 | 2 (1 from cortex) |

LDA = linear discriminant analysis; KNN = k-nearest neighbor; MLP = multilayer perceptron neural network; AUC = Area under the Receiver Operating Characteristic Curve.

### Ex-vivo IHC confirms microglia uptake as the mechanism for NCDx as the disease biomarker

Having demonstrated strong ability to separate disease from control, we next assessed the specificity of probe binding to microglia in *vivo*. To make this assessment, we began by establishing the relationship between NCDx signal as confirmed using the intrinsic fluorescent Dil dye (Fig. **2**) and microglia using an antibody for ionized calcium-binding adaptor molecule 1 (IBA-1). Tissue sections from TG mice stained with the antibody showed focal co-localization of NCDx with IBA-1 reactivity (Fig. **5A**). 3D Imaris images generated from a composite image showed that the NCDx signal was in the cytosolic compartment of the IBA-1 reactive cells (Fig. **5A** top row). Tissue sections from the WT controls showed IBA-1 reactivity but no organized NCDx signal, suggesting no significant uptake of NCDx by microglia in the WT brain, consistent with the *in vivo* MRI data. Whole-scan images of the olfactory bulb (fig. **S7 - S14**) revealed more detailed and consistent with the *in vivo* MRI radiomics findings.

**Fig. 5.**
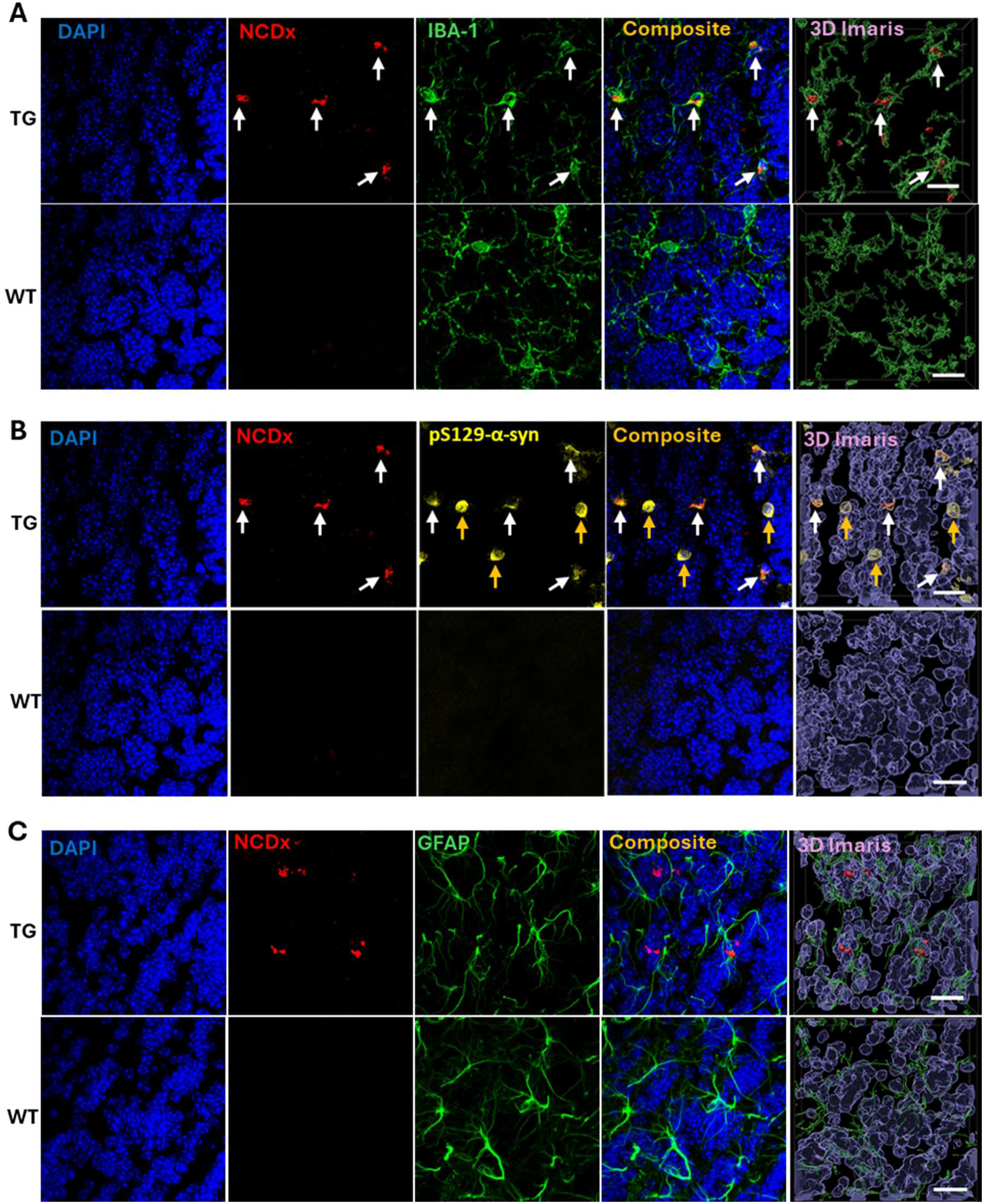
*Ex vivo* IHC demonstrating in *vivo* accumulation of NCDx is driven primarily by microglia uptake. Brain tissue sections from treated TG mice show (**A**) NCDx signal in the cytosolic compartment of IBA-1 reactive cells; (**B**) NCDx accumulation correlates with the presence of Lewy pathology; (**C**) NCDx signal does not show direct overlap or co-localization with GFAP reactivity. Scale bar = 20 µm.

Next, we assessed the relationship between NCDx signal and Lewy pathology staining the tissue with a phosphoserine 129 (pS129–α-syn, pathologic α-syn) antibody. Well organized structures without the corresponding NCDx signal (orange arrows) attributed to Lewy bodies were also observed in tissue sections from TG mice (Fig. **5B**, top row). Tissue sections from WT mice stained with the same antibody showed neither signal from NCDx nor signal from pS129–α-syn antibody (bottom row). These observations suggest that the uptake of NCDx is correlated with the presence of Lewy pathology.

Finally, to confirm that in *vivo* NCDx uptake was specific to CSF1-R expressing cells, contiguous sections from both TG and WT mice were further stained with anti-GFAP antibody to assess any relationship between NCDx and astrocytes. TG tissue sections showed both NCDx signal and GFAP reactivity, but composite images did not show any meaningful overlap between the two signals, even in 3D Imaris images (Fig. **5C** top row). Tissue sections from WT controls showed GFAP reactivity but no significant NCDx signal as expected (Fig. **5C** bottom row), suggesting that astrocytes, which are not known to express the CSF1-R receptor, do not internalize NCDx, consistent with earlier observations in the in *vitro* cell uptake studies.

Quantitative colocalization analysis of NCDx with cell-specific markers in tissue from TG mice injected with NCDx was performed using ImarisColoc (Figure **6**). The Pearson correlation coefficient (PCC) was 0.50 for NCDx–IBA-1 signals (Fig. **6E**) and −0.04 for NCDx–GFAP signals (Fig. **6J**), indicating substantial colocalization with microglia but not astrocytes. Together with the cell uptake data, these results suggest that NCDx preferentially accumulates in microglial cells. Whole-scan images of the olfactory bulb revealed more detailed and consistent findings (Fig. **S11** and **S12**).

**Fig. 6.**
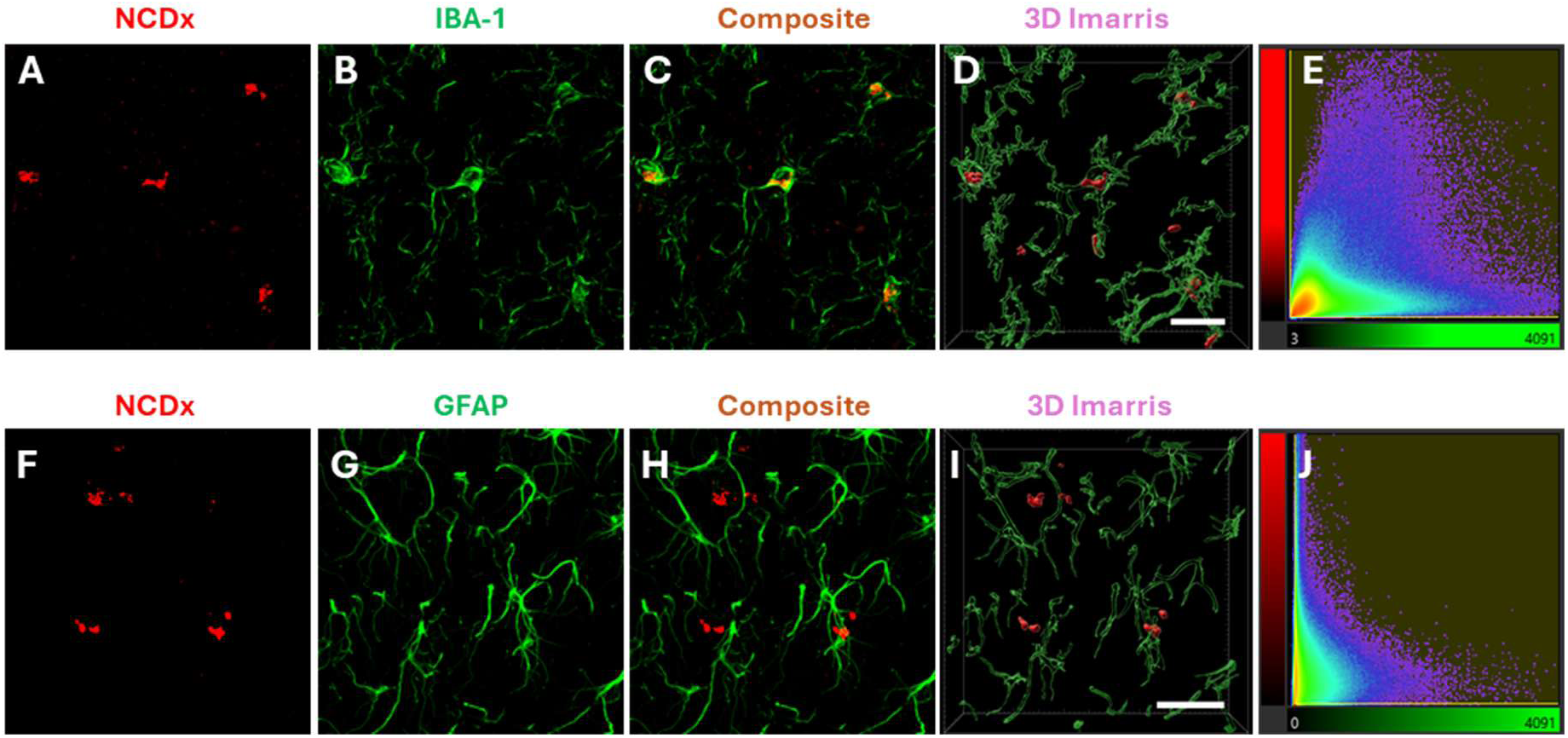
Colocalization analysis of the NCDx and antibody signals. (**A**) Fluorescence due to accumulation of NCDx in the cytosolic compartment of IBA-1 reactive cells; (**B**) Fluorescence from antibody IBA-1; (**C**) Merged NCDx and IBA-1 signals; (**D**) ImarisColoc 3D image of fluorescence intensities within the colocalized volume; (**E**) Scatter plot of pixels within the colocalized volume; (**F**) Fluorescence due to accumulation of NCDx in IBA-1 reactive cells or other macrophage cells; (**G**) Fluorescence from antibody GFAP; (**H**) Merged NCDx and GFAP signals; (**I**) ImarisColoc 3D image of fluorescence intensities within the colocalized volume; **(J**) Scatter plot of pixels within the colocalized volume. Scale bar = 20 µm

## DISCUSSION

Using a combination of *in silico* modeling, *in vitro* cell uptake studies, and *in vivo* imaging of a mouse model of PD, we showed that PEGylated Caflanone, the targeting moiety on **NCDx**, has the potential to serve as a sensitive and specific biomarker microglia-mediated neuroinflammation in PD. **NCDx** engages the extracellular Ig domains of CSF1-R, a receptor selectively enriched on activated microglia allowing the nanoparticles to be rapidly internalized by microglial cell lines in a dose-dependent manner. Importantly, following intravenous administration in *vivo*, AI-assisted image analysis of T1-weighted MR scans enabled noninvasive separation of Parkinsonian mice from controls, reflecting the heightened microglial activation characteristic of chronic neuroinflammation and Lewy pathology distribution in this model.

Most small-molecule CSF1-R binding ligands (inhibitors) in advanced stages of clinical development or approved by the FDA (39–42), bind the intracellular tyrosine kinase domain of the protein, blocking downstream signaling but offering no capacity for cell-type–specific targeting or extracellular discrimination. In contrast, our ligand engages the extracellular Ig-like domains of CSF1-R, the same structural region used by endogenous cytokines CSF-1 and IL-34 to initiate receptor dimerization and activation. This mode of engagement is significant for two reasons. First, extracellular binding enables selective recognition of CSF1R-expressing myeloid cells, particularly chronically activated microglia without requiring intracellular access or kinase inhibition, thereby allowing nanoparticle delivery of the imaging payload without globally suppressing CSF1-R signaling. Second, extracellular targeting leverages the receptor’s natural ligand-binding architecture, which is structurally well-defined and conserved across species, enabling high-affinity, surface-restricted interactions that are ideal for receptor-mediated endocytosis. This creates a mechanistically distinct and translationally powerful entry point, cell-specific delivery of diagnostic and potentially therapeutic payloads.

The ability of NCDx to enable noninvasive separation of parkinsonian mice from controls through microglial-targeted mMRI is highly significant for the Parkinson’s disease research community because it directly addresses a major unmet need: a specific, *in vivo* biomarker of neuroinflammation that tracks early disease biology rather than downstream neurodegeneration (^43,44^). Microglia activation and CSF1-R upregulation (^45,46^) are increasingly recognized as central features of PD pathogenesis, emerging early and potentially driving α-synuclein aggregation and nigrostriatal injury, yet current clinical imaging has relied on TSPO ligands with limited specificity and interpretability (^47,48^). Demonstrating that a CSF1R-targeted nanoparticle can noninvasively distinguish parkinsonian mice from controls shows that chronic neuroinflammation is not only detectable but also quantifiable and spatially resolvable, opening the door to earlier diagnosis, mechanistically grounded patient stratification, and dynamic monitoring of microglia-directed therapies. Importantly, because dysregulated microglial activation is a shared, early, and mechanistically causal feature across other neurodegenerative disorders such as Alzheimer’s disease, amyotrophic lateral sclerosis, frontotemporal dementia, multiple sclerosis, and traumatic brain injury, a CSF1R-specific MRI agent will enable unprecedented opportunities for early diagnosis, biologically grounded patient stratification, and real-time monitoring of neuroimmune-directed therapies, if successful developed.

In conclusion, our findings establish a mechanistically targeted, microglia-engaging nanoparticle platform capable of detecting and reporting chronic neuroinflammation in *vivo* with cellular specificity. By demonstrating selective binding to the extracellular domain of CSF1R, rapid receptor-mediated uptake by microglia, and MRI-based discrimination of parkinsonian mice from controls, this work provides a new noninvasive window into the inflammatory states that shape Parkinson’s disease progression. The concordance between in *vivo* imaging and ex *vivo* IHC confirms that the agent faithfully reports microglial activation rather than nonspecific tissue changes, addressing a long-standing gap in neuroinflammation biomarkers. More broadly, this platform offers a scalable foundation for early diagnosis, longitudinal monitoring, and targeted therapeutic delivery in neurodegenerative diseases where microglial dysfunction is central.

## LIMITATIONS

Although these findings establish a promising platform for detecting microglial activation in *vivo*, some limitations warrant consideration. First, while the nanoparticle demonstrated selective engagement of CSF1-R-expressing microglia, the study did not resolve how uptake varies across microglial subtypes or disease-stage, which may influence sensitivity and specificity in more heterogeneous models. Second, the imaging experiments were performed in a single Parkinsonian mouse model, thus it remains unclear how well the agent will generalize across other α-synuclein-driven or toxin-based models that exhibit distinct inflammatory signatures. Finally, while ex-*vivo* IHC confirmed microglial uptake, we did not address long-term biodistribution, potential off-target interactions, or the fate of cells following nanoparticle internalization. Future work in these areas will be essential for advancing this platform toward clinical translation as a noninvasive biomarker of neuroinflammation in Parkinson’s disease.

## MATERIALS AND METHODS

### Computational modeling studies

The crystal structure of extracellular CSF-1R: CSF-1complex (PDB 4WRM^1^, resolution 6.85Å) was preprocessed with Schrödinger Maestro 14.1. Preprocessing using the protein preparation wizard of Maestro involved filling in any missing side chains and loops, adding missing hydrogens, enumerating bond orders, determining protonation states for het groups at pH 7.4 ± 2.0, optimizing H-bonds, followed by restrained minimization and deletion of waters molecules that are not in close proximity to het groups (^49^).The quality of the prepared protein was assessed with a protein reliability report which showed no red flags indicating the prepared proteins was reliable for computational modeling.

PEGylated Caflanone was preprocessed using Maestro’s LigPrep tool. Briefly, we converted the 2D structure of the ligand to multiple 3D structures, added hydrogen atoms, determined partial charges of atoms, and generated multiple conformers and ionizations states at pH 7.4 ± 2.0 of the ligands (^49^).

We used the SiteMap Maestro tool to explore possible druggable binding sites across CSF1-R_D1-D5_. We opted for SiteMap to report up to five top sites and required the top sites to have at least 15 site points per site. SiteScores (SScore) and druggability Scores (DScore) were generated and used to rank the sites (^50,51^). A receptor grid was generated using the site points reported from the potential druggable sites identified by SiteMap. The coordinates of the receptor grid were expanded to X, Y, Z of 15 Å at all sites for docking PEGylated Caflanone due to the larger size of the molecule. All other default parameters were retained. Glide was used to dock the prepared PEGylated Caflanone ligand conformers using extra precision (XP) docking. The model was set to prioritize intramolecular H-bonds and enhance the planarity of conjugated π groups. Glide scores that were generated were used to rank the best pose of the ligand when docked within the top ranked selected potential binding sites (^52,53^).

### Nanoparticle formulation

We synthesized and characterized Caflanone-PEG (3400)-DSPE conjugate, the targeting component of the formulation as described in supplemental methods section. NCDx and the control nontargeted liposome formulation were prepared using standard hydration/extrusion protocol. Briefly, a lipid mixture containing HSPC, DSPE-mPEG(2000), cholesterol, Gd(III)DSPE-DOTA, and the Caflanone-DSPE_3.4k_-conjugate (replaced with DSPE-mPEG(2000) in the nontargeted formulation) at a molar ratio of 32:2.5:40:25:0.5 in ethanol (600 μL) was stirred at 60–65 °C until all solids dissolved to form a clear solution. 50 μL of DiLC dye dissolved in ethanol (1 mg in 200 μL) was added to the lipid solution and stirred for 5 mins. 12 mL histidine-buffered saline (HBS) (10 mM histidine, 140 mM NaCl, ∼pH 7.6) was added to the lipid solution and stirred at 60–65 °C for 45 minutes. The hydrated lipid solution was extruded sequentially through 400 nm (5 passes) and 200 nm (8 passes) Nuclepore membranes at 60–65 °C using a high-pressure extruder (Northern Lipids, Vancouver, BC, Canada) to form liposomes. The liposomal suspension was dialyzed against HBS using 300 kDa molecular weight cutoff membranes (Spectrum Laboratories Inc., CA, USA) to remove any unencapsulated lipids and ethanol. The nanoparticle size was determined by dynamic light scattering (DLS), and the concentrations of Gd(III) and total lipids in the final formulation were determined by inductively coupled plasma‒optical emission spectrometry (ICP‒OES).

### Cell uptake studies

SIM-A9 cells (ATCC, Manassas, VA, U.S.A.), RAW264.7 (ATCC, Manassas, VA, U.S.A.), mouse primary astrocytes (isolated and cultured from C57Bl/6 mice), and human primary astrocytes (ScienCell, CA, USA) cultured in Dulbecco’s Modified Eagle Medium: Nutrient Mixture F12 (DMEM/F12, Thermo Fisher Scientific, Waltham, MA, U.S.A.), separately. The human microglial clone 3 (HMC-3) cell line (ATCC, Manassas, VA, USA) cultured in Eagle’s minimum essential media (EMEM, ATCC). All cell lines were supplemented with 10% fetal bovine serum (FBS, Thermo Fisher Scientific for SIM-A9 and Sigma‒Aldrich, St. Louis, MO, USA for HMC-3) and 100 units/mL (U/mL) penicillin/streptomycin (Invitrogen, Carlsbad, CA, USA) and incubated in a humidified atmosphere (5% CO_2_) at 37 °C.

HMC-3 (2*10^4^ cells/chamber) were seeded on 8 chamber culture slides (Falcon™ 354118) and allowed to adhere overnight. NCDx (80 µM total lipid) were added to the cells. After 2 hours of incubation, the supernatant was removed by pipetting, and the cells were washed with PBS three times. The cells were fixed in 4% paraformaldehyde for 15 min and then washed with PBS three times. They were then permeabilized with 0.1% Triton X-100 for 10 min and washed with PBS three times. The cells were then blocked with 3% normal goat serum for 30 min, the supernatant removed by pipetting, and incubated with phalloidin 488 (1:300, ab176753, Abcam) for 1 hour. This was followed by another wash with PBS and then incubation with Hoechst (33342, Thermo Fisher Scientific) at room temperature for 7 minutes. Finally, the cells were washed with PBS three times, coverslipped, and imaged under an Olympus IX81 microscope using standard excitation/emission filters. All other cell lines were cultured, stained, and imaged using the same procedures.

### Activation study with CSF1

Initially, 5000 cells per well were seeded in PerkinElmer Pheno plate 96-well. Cells were allowed to get attached for overnight at 37 °C incubator with 5% CO_2_ and 90% relative humidity. Cells were then exposed to 25 ng/ml of colony stimulator factor 1 (CSF1) for 24 hours (^54^). Cells were treated with 80 µM NCDx or non-targeted nanoparticles for 2 hours for cell uptake study and up to 2, 4 or 22 hours for kinetics study at different timepoints. Cells were washed twice with 1XPBS and then fix-perm (Fixation-permeabilization with 10% formalin containing 0.2% triton X-100) for 10 minutes, washed thrice. For cytoskeleton staining, CellMask Actin-488 (ThermoFisher, USA) was used, and nucleus was stained with Hoechst dye. Plate was imaged with Olympus IX81 microscope at 40X objective. Formula used to calculate ET_50_= T0+0.5*(T_max_-T_min_).

### Caflanone blocking studies

SIM-A9 (2*10^4^ cells/chamber) were seeded on 8 chamber culture slides (Falcon™ 354118) and allowed to adhere overnight. NCDx (40 µM total lipid) and different concentrations of free ligand (Caflanone) from 0 µM to 50 µM were added to the cells at the same time. After 2 hours of incubation, the supernatant was removed by pipetting, and the cells were washed with PBS three times. The cells were fixed in 4% paraformaldehyde for 15 min and then washed with PBS three times. They were then permeabilized with 0.1% Triton X-100 for 10 min and washed with PBS three times. The cells were then blocked with 3% normal goat serum for 30 min, the supernatant removed by pipetting, and incubated with phalloidin 488 (1:300, ab176753, Abcam) for 1 hour. This was followed by another wash with PBS and then incubation with Hoechst (33342, ThermoFisher, USA) at room temperature for 7 minutes. Finally, the cells were washed with PBS three times, coverslipped, and imaged under an Olympus IX81 microscope using standard excitation/emission filters.

### Animal studies

A53T transgenic mice (^55^) (N = 8, 12- to 14-month-olds) and age-matched wildtype controls (N = 8) were used for this study. All animal studies reported in this paper were conducted under study-specific protocols approved by the Institutional Animal Care and Use Committee (IACUC) at Baylor College of Medicine.

### In vivo MRI

*In vivo* MRI studies were performed on a 1T permanent magnet (M2 system, Aspect Imaging, Shoham, Israel). For each imaging experiment, animals were sedated using 2-3% isoflurane and then placed on a custom fabricated bed with an integrated face-cone for continuous anesthesia delivery by inhalation (1.5% isoflurane) for the duration of the experiment. Respiration rate was monitored by a pneumatically controlled pressure pad placed underneath the animal’s abdomen. To establish a baseline (pre-contrast), each mouse was subjected to an axial T1-weighted spin echo (T1w-SE) sequence with the following parameters: TR = 600 ms, TE = 11.5 ms, matrix = 192x192, FOV = 30 mm, slices = 16, slice thickness = 1.2 mm, NEX = 2. Following the pre-contrast scans, mice administered NCDx intravenously via tail vein at a dose of 0.15 mmol Gd(III)/kg and returned to their cages. Four days later, when the contrast had cleared systemic circulation, they were subjected to the same scan protocol to obtain post-contrast imaging data. Following this scan, each mouse was euthanized, perfused with PBS and 4% formalin, and the brains were excised and stored in 30% sucrose solution until sectioning for *ex-vivo* immunohistochemical analysis (see below).

The raw imaging data was converted to dicoms and analyzed in Osirix (version 5.8.5, 64-bit). Region of interests (ROIs) were drawn in the cortex (bregma 0 mm) and olfactory bulbs (bregma 4-5 mm) of all animals in the pre-contrast and post-contrast scans. Each region was taken in three contiguous 1.2 mm slices. All images were analyzed on the same MR signal intensity scale (1500-4500 a.u.). Differences in subjects were analyzed as relative differences in signal change, which was calculated as the percentage increase in signal in the post-contrast scan relative to the baseline, pre-contrast scan (% Signal Change). All statistical analysis was performed using a non-parametric Wilcoxon rank-sum test (MATLAB, version 2025).

### Machine learning - Radiomics

Post-contrast MRIs were first processed by registering a high-resolution brain atlas, annotated with 27 brain regions. This was followed by automatic extraction of cerebral cortex and olfactory bulb areas, using 3D Slicer. RFs were extracted using the Python package PyRadiomics (^56^), with and without applying filters (wavelet and Laplacian of Gaussian), following recommendations from the International Biomarker Standardization Initiative (IBSI) (^57^), which identifies a subset of reproducible RFs suitable for research. RFs from cerebral cortex and/or olfactory bulb were examined in parallel. Although the same scanner and imaging protocol were used and images were acquired within a similar time frame, to confirm the robustness of our findings, we performed a parallel analysis in which RFs were extracted from three datasets: (i) original raw images, (ii) images resampled to isotropic resolution (0.15 × 0.15 × 0.15 mm), and (iii) isotropic images with voxel intensities minmax normalized.

A standard ML workflow was applied to classify Tg and Wt mice. Briefly, in each classification task, the dataset was split into 80% training/validation (6 Tg, 6 Wt) and 20% testing (2 TG, 2 WT). Feature selection in the training set involved removing highly correlated features (Spearman correlation > 0.8) and selecting top-ranked features based on importance scores calculated using Neighborhood Component Analysis. Various classifiers were evaluated, including logistic regression, k-nearest neighbor (KNN), linear discriminant analysis (LDA), support vector machine (SVM), multilayer perceptron neural network (MLP), random forest, and ensemble models. Hyperparameters were optimized during 3-fold cross-validation on the training/validation set. The final classifier was selected based on validation performance. The same training and testing sets were used across all parallel analyses with respect to feature types and image processing strategies. All analyses were performed using Python ver 3.12.2and image segmentation was done in 3D slicer ver 5.2.2.

### *Ex-vivo* immunohistochemical analysis

Brains were embedded in Tissue-Tek O.C.T. and kept in liquid nitrogen for 30 minutes. The embedded tissue was sliced into 30 μm thick sections with Lecia Biosystems Cryostats at -20 °C. Tissue sections were washed twice with PBS, followed by antigen retrieval with citric acid buffer (pH ∼ 6.2)/microwave for 15 minutes. The sections were then washed with PBS three times and permeabilized with 0.2% Triton-X 100 for 10 minutes, followed by washing with PBS two times. The tissue was incubated with 3% normal goat serum at room temperature for one hour, followed by incubation with primary antibodies, an anti-alpha-synuclein (phospho S129) antibody [EP1536Y] (ab51253), Abcam) (anti-rabbit, 1:500 in 1% goat serum) and Anti-Iba1/AIF1 Antibody [anti-Mouse, 1:500 in 1% goat serum, (EMD Millipore, MABN92)] overnight at 4 °C. The tissue was washed with PBS three times after overnight incubation and then incubated for two hours at room temperature with Alexa Fluor 488 (anti-mouse) and Alexa Fluor 647 (anti-rabbit) secondary antibody (1:500 in PBS). Following two additional PBS washes, the tissue was incubated with Hoechst (33342, Thermo Fisher Scientific) at room temperature for 7 minutes. The sections were washed with PBS three times, coverslipped, and imaged under an Olympus IX81 microscope using standard excitation/emission filters.

### Statistical analysis

Statistical analyses for the *in vitro* cell experiments that followed a normal distribution were performed using one-way ANOVA with pairwise comparisons adjusted using Fisher’s least significant difference (LSD) test. Statistical analyses of *in vivo* experimental data that did not follow a normal distribution were performed using the Kruskal‒Wallis test with Bonferroni-adjusted pairwise comparisons.

## Supplementary Materials

The PDF file includes

Materials and Methods (chemical synthesis and characterization of synthetic intermediates and final products)

Figs. S1 to S14

^1^H NMR, ^13^NMR, and MALDI spectra.

**Other Supplementary Material for this manuscript includes the following:**

Movies M1 to M4

## Acknowledgments

This manuscript was prepared with the assistance of a science writer, Ariel M Lyons-Warren, MD PhD who is supported by Texas Children’s Hospital’s Research Vision Program.

## Author contributions

Conceptualization: NT, ET.

Methodology: XS, AB, T-ER, EN, NT, ET.

Investigation: XS, AB, T-ER, EN, SM, NT, ET.

Visualization:

Funding acquisition: ET

Project administration: ET

Supervision: ET.

Writing – original draft: XS, AB, T-ER, EN, SM, ET.

Writing – review & editing: XS, AB, T-ER, EN, SM, JC, AA, HL, NT, ET.

## REFERENCES

1. Li, M., Ye, X., Huang, Z., Ye, L. & Chen, C. Global burden of Parkinson’s disease from 1990 to 2021: a population-based study. BMJ Open 15, e095610 (2025).

2. Statistics | Parkinson’s Foundation. https://www.parkinson.org/understanding-parkinsons/statistics.

3. Wüllner, U., et al. The heterogeneity of Parkinson’s disease. J Neural Transm (Vienna) 130, 827–838 (2023).

4. Morris, H. R., Spillantini, M. G., Sue, C. M. & Williams-Gray, C. H. The pathogenesis of Parkinson’s disease. The Lancet 403, 293–304 (2024).

5. Tolosa, E., Garrido, A., Scholz, S. W. & Poewe, W. Challenges in the diagnosis of Parkinson’s disease. Lancet neurology 20, 385–397 (2021).

6. Isik, S., Yeman Kiyak, B., Akbayir, R., Seyhali, R. & Arpaci, T. Microglia Mediated Neuroinflammation in Parkinson’s Disease. Cells 12, 1012 (2023).

7. McGeer, P. L., Itagaki, S., Boyes, B. E. & McGeer, E. G. Reactive microglia are positive for HLA-DR in the substantia nigra of Parkinson’s and Alzheimer’s disease brains. Neurology 38, 1285–1291 (1988).

8. Imamura, K., et al. Distribution of major histocompatibility complex class II-positive microglia and cytokine profile of Parkinson’s disease brains. Acta Neuropathol 106, 518–526 (2003).

9. Castillo-Rangel, C., et al. Neuroinflammation in Parkinson’s Disease: From Gene to Clinic: A Systematic Review. International journal of molecular sciences 24, 5792- (2023).

10. Eser, P., Kocabicak, E., Bekar, A. & Temel, Y. The interplay between neuroinflammatory pathways and Parkinson’s disease. Experimental neurology 372, 114644–114644 (2024).

11. Wüllner, U., et al. The heterogeneity of Parkinson’s disease. Journal of Neural Transmission 130, 827–838 (2023).

12. Liu, T.-W., Chen, C.-M. & Chang, K.-H. Biomarker of Neuroinflammation in Parkinson’s Disease. International journal of molecular sciences 23, 4148- (2022).

13. Braak, H., Ghebremedhin, E., Rüb, U., Bratzke, H. & Del Tredici, K. Stages in the development of Parkinson’s disease-related pathology. Cell Tissue Res 318, 121–134 (2004).

14. Li, T., Qiu, T., Jiang, F., Cai, H. & Le, W. Microglia in the crosstalk between peripheral and central nervous systems in Parkinson’s disease. Transl Neurodegener 14, 66 (2025).

15. Lind-Holm Mogensen, F., Seibler, P., Grünewald, A. & Michelucci, A. Microglial dynamics and neuroinflammation in prodromal and early Parkinson’s disease. Journal of neuroinflammation 22, 136–17 (2025).

16. Oh, S. J., et al. Evaluation of the Neuroprotective Effect of Microglial Depletion by CSF-1R Inhibition in a Parkinson’s Animal Model. Molecular imaging and biology 22, 1031–1042 (2020).

17. Mills, K. A., et al. Exploring [11C]CPPC as a CSF1R-targeted PET imaging marker for early Parkinson’s disease severity. The Journal of clinical investigation 135, (2025).

18. Isik, S., Yeman Kiyak, B., Akbayir, R., Seyhali, R. & Arpaci, T. Microglia Mediated Neuroinflammation in Parkinson’s Disease. *Cells (Basel*, Switzerland) 12, 1012- (2023).

19. Roodveldt, C., et al. The immune system in Parkinson’s disease: what we know so far. Brain (London, England : 1878) 147, 3306–3324 (2024).

20. Vivash, L. & O’Brien, T. J. Imaging Microglial Activation with TSPO PET: Lighting Up Neurologic Diseases? Journal of Nuclear Medicine 57, 165–168 (2016).

21. Lavisse, S., et al. Increased microglial activation in patients with Parkinson disease using [18F]-DPA714 TSPO PET imaging. Parkinsonism & related disorders 82, 29–36 (2021).

22. Kreisl, W. C., et al. A genetic polymorphism for translocator protein 18 kDa affects both in vitro and in vivo radioligand binding in human brain to this putative biomarker of neuroinflammation. J Cereb Blood Flow Metab 33, 53–58 (2013).

23. Lee, N., Choi, J. Y. & Ryu, Y. H. The development status of PET radiotracers for evaluating neuroinflammation. Nucl Med Mol Imaging 58, 160–176 (2024).

24. Uzuegbunam, B. C., Rummel, C., Librizzi, D., Culmsee, C. & Hooshyar Yousefi, B. Radiotracers for Imaging of Inflammatory Biomarkers TSPO and COX-2 in the Brain and in the Periphery. International journal of molecular sciences 24, 17419- (2023).

25. van der Wildt, B., et al. Discovery of a CSF-1R inhibitor and PET tracer for imaging of microglia and macrophages in the brain. Nuclear medicine and biology 114-115, 99–107 (2022).

26. Zha, X., Hui, W., Cheng, D., Shi, H. & Luo, Z. Targeting colony-stimulating factor 1 receptor: From therapeutic drugs to diagnostic radiotracers. iRadiology (Online) 2, 35–65 (2024).

27. Salarian, M., et al. Evaluation of [18F]JNJ-CSF1R-1 as a Positron Emission Tomography Ligand Targeting Colony-Stimulating Factor 1 Receptor. Molecular imaging and biology 27, 163–172 (2025).

28. Tanifum, E. A., et al. Intravenous delivery of targeted liposomes to amyloid-β pathology in APP/PSEN1 transgenic mice. PLoS One 7, e48515 (2012).

29. Tanifum, E. A., et al. A Novel Liposomal Nanoparticle for the Imaging of Amyloid Plaque by Magnetic Resonance Imaging. J Alzheimers Dis 52, 731–745 (2016).

30. Badachhape, A. A., et al. Pre-clinical dose-ranging efficacy, pharmacokinetics, tissue biodistribution, and toxicity of a targeted contrast agent for MRI of amyloid deposition in Alzheimer’s disease. Sci Rep 10, 16185 (2020).

31. Sun, X., et al. A dual target molecular magnetic resonance imaging probe for noninvasive profiling of pathologic alpha-synuclein and microgliosis in a mouse model of Parkinson’s disease. Frontiers in neuroscience 18, 1428736- (2024).

32. Lowe, H. & Toyang, N. J. Therapeutic agents containing cannabis flavonoid derivatives targeting kinases, sirtuins and oncogenic agents for the treatment of multiple myeloma, lymphoma, and leukemia. (2021).

33. Hu, B., et al. Insights Into the Role of CSF1R in the Central Nervous System and Neurological Disorders. Front. Aging Neurosci. 13, 789834 (2021).

34. Alzyoud, L., Bryce, R. A., Al Sorkhy, M., Atatreh, N. & Ghattas, M. A. Structure-based assessment and druggability classification of protein–protein interaction sites. Scientific reports 12, 7975–18 (2022).

35. Giasson, B. I., Duda, J. E., Quinn, S. M., Zhang, B. & Trojanowski, J. Q. Neuronal ␣-Synucleinopathy with Severe Movement Disorder in Mice Expressing A53T Human ␣-Synuclein.

36. Van Griethuysen, J. J. M., et al. Computational Radiomics System to Decode the Radiographic Phenotype. Cancer Research 77, e104–e107 (2017).

37. Whybra, P., et al. The Image Biomarker Standardization Initiative: Standardized Convolutional Filters for Reproducible Radiomics and Enhanced Clinical Insights. Radiology 310, e231319 (2024).

38. Hatt, M., et al. Radiomics: Data Are Also Images. Journal of Nuclear Medicine 60, 38S–44S (2019).

39. Chompunud Na Ayudhya, C., Graidist, P. & Tipmanee, V. Role of CSF1R 550th-tryptophan in kusunokinin and CSF1R inhibitor binding and ligand-induced structural effect. Sci Rep 14, 12531 (2024).

40. Fujiwara, T., et al. CSF1/CSF1R Signaling Inhibitor Pexidartinib (PLX3397) Reprograms Tumor-Associated Macrophages and Stimulates T-cell Infiltration in the Sarcoma Microenvironment. Mol Cancer Ther 20, 1388–1399 (2021).

41. Kane, J. L., et al. Identification of Selective Imidazopyridine CSF1R Inhibitors. ACS Med Chem Lett 15, 722–730 (2024).

42. Peng, L., Fang, Z., Zhang, W., Rao, G.-W. & Zheng, Q. Recent advances in colony stimulating factor-1 receptor (CSF1R) inhibitors. Biochemical Pharmacology 242, 117187 (2025).

43. Chen, K., Wang, H., Ilyas, I., Mahmood, A. & Hou, L. Microglia and Astrocytes Dysfunction and Key Neuroinflammation-Based Biomarkers in Parkinson’s Disease. Brain Sciences 13, 634 (2023).

44. Lind-Holm Mogensen, F., Seibler, P., Grünewald, A. & Michelucci, A. Microglial dynamics and neuroinflammation in prodromal and early Parkinson’s disease. J Neuroinflammation 22, 136 (2025).

45. Lee, S. J., Wang, C. & Hooker, J. Microglia matters: visualizing the immune battle in Parkinson’s disease. J Clin Invest 135, e192919 (2025).

46. Mills, K. A., et al. Exploring [^11^C]CPPC as a CSF1R-targeted PET imaging marker for early Parkinson’s disease severity. J Clin Invest 135, (2025).

47. Owen, D. R. J., et al. Mixed-Affinity Binding in Humans with 18-kDa Translocator Protein Ligands. Journal of Nuclear Medicine 52, 24–32 (2011).

48. Vivash, L. & O’Brien, T. J. Imaging Microglial Activation with TSPO PET: Lighting Up Neurologic Diseases? Journal of Nuclear Medicine 57, 165–168 (2016).

49. Madhavi Sastry, G., Adzhigirey, M., Day, T., Annabhimoju, R. & Sherman, W. Protein and ligand preparation: parameters, protocols, and influence on virtual screening enrichments. J Comput Aided Mol Des 27, 221–234 (2013).

50. Halgren, T. New method for fast and accurate binding-site identification and analysis. Chem Biol Drug Des 69, 146–148 (2007).

51. Halgren, T. A. Identifying and characterizing binding sites and assessing druggability. J Chem Inf Model 49, 377–389 (2009).

52. Liang, J., et al. Site mapping and small molecule blind docking reveal a possible target site on the SARS-CoV-2 main protease dimer interface. Computational Biology and Chemistry 89, 107372 (2020).

53. Ngwa, W., et al. Potential of Flavonoid-Inspired Phytomedicines against COVID-19. Molecules 25, 2707 (2020).

54. Morandi, A., Barbetti, V., Riverso, M., Dello Sbarba, P. & Rovida, E. The Colony-Stimulating Factor-1 (CSF-1) Receptor Sustains ERK1/2 Activation and Proliferation in Breast Cancer Cell Lines. PLoS ONE 6, e27450 (2011).

55. Lee, M. K., et al. Human α-synuclein-harboring familial Parkinson’s disease-linked Ala-53 → Thr mutation causes neurodegenerative disease with α-synuclein aggregation in transgenic mice. Proceedings of the National Academy of Sciences 99, 8968–8973 (2002).

56. van Griethuysen, J. J. M., et al. Computational Radiomics System to Decode the Radiographic Phenotype. Cancer Res 77, e104–e107 (2017).

57. Whybra, P., et al. The Image Biomarker Standardization Initiative: Standardized Convolutional Filters for Reproducible Radiomics and Enhanced Clinical Insights. Radiology 310, e231319 (2024).

