## Supplementary Materials for "Microglia-Specific Molecular Magnetic Resonance Imaging Probe Enables Noninvasive Separation of Parkinsonian Mice from Controls"

**The PDF file includes:**

Materials and Methods

Figs. S1 to S14

NMR spectrum SN1 to SN12

**Other Supplementary Material for this manuscript includes the following:**

Movies S1 to S4

### MATERIALS AND METHODS

#### Materials

Unless otherwise noted, all reagents and solvents were obtained from commercial sources including Sigma-Aldrich, TCI, and Acros Organics and used without further purification. Thin layer chromatography (TLC) was performed on silica gel 60 F254 plates from EMD Chemical Inc. and components were visualized by ultraviolet light (254 nm) and/or phosphomolybdic acid, 20 wt% solution in ethanol. SiliFlash silica gel (230–400 mesh) was used for all column chromatography.

#### Measurements

Proton nuclear magnetic resonances (1H NMR) were recorded at 600 MHz or 500 MHz on Bruker 600 or 500 NMR spectrometers. Carbon nuclear magnetic resonances (13C NMR) were recorded at 75 MHz or 125 MHz on a Bruker 300 or 500 NMR spectrometers respectively. Chemical shifts are reported in parts per million (ppm) from an internal standard chloroform (7.26 ppm), or dimethylsulfoxide (2.50 ppm) for 1H NMR; and from an internal standard of either residual chloroform (77.00 ppm), or dimethylsulfoxide (39.52 ppm) for 13C NMR. NMR peak multiplicities are denoted as follows: s (singlet), d (doublet), t (triplet), q (quartet), dd (doublet of doublet), td (doublet of triplet), dt (triplet of doublet), and m (multiplet). Coupling constants (J) are given in hertz (Hz). High resolution mass spectra (HRMS) were obtained from The Ohio State University Mass Spectrometry and Proteomics Facility. The mean particle size of liposomes was determined using a Dynamic Light Scattering (DLS) instrument (Brookhaven Instruments Corp., Holtsville, NY, USA). Formulation elemental analysis was determined using Inductively Coupled Plasma Optical Emission Spectroscopy (ICP-OES). Magnetic Resonance Imaging (MRI) was performed on a 1 T permanent magnet scanner (M2 system, Aspect Imaging, Shoham, Israel). All confocal microscopy images were captured from an Olympus IX81 microscope, and the images were processed and analyzed using Fiji-ImageJ software.

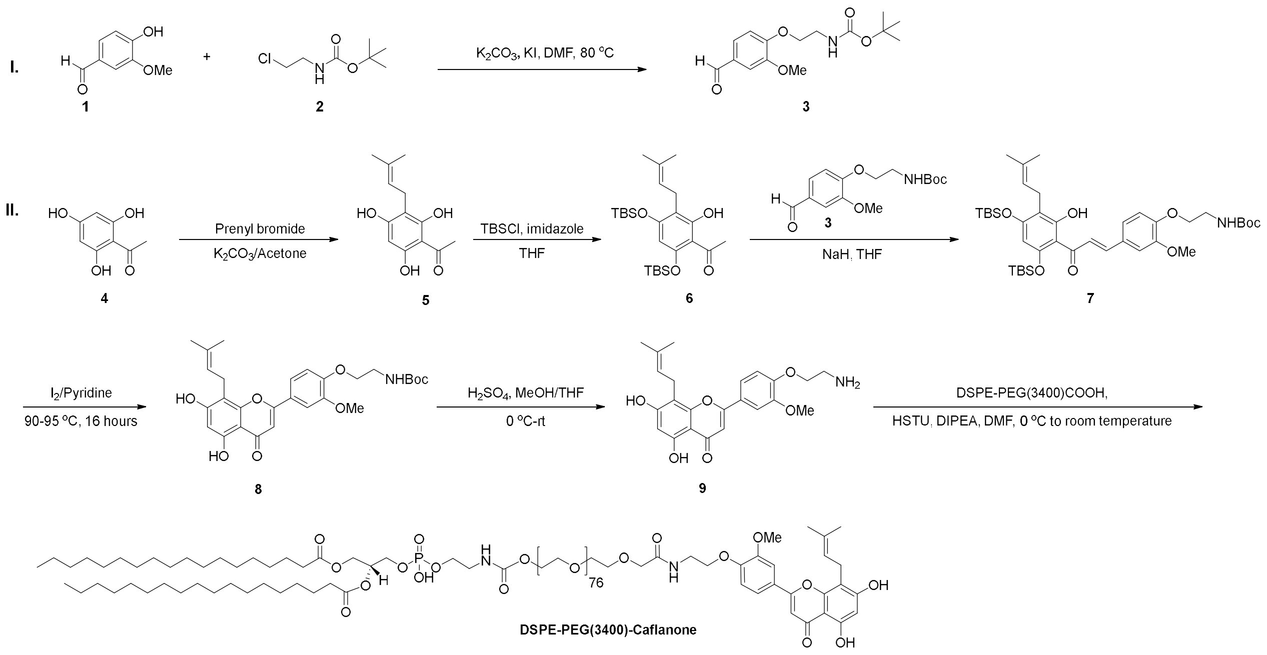

**Scheme 1.** Synthetic route to Caflanone-PEG(3400)-DSPE conjugate.

### Synthesis of compound 3

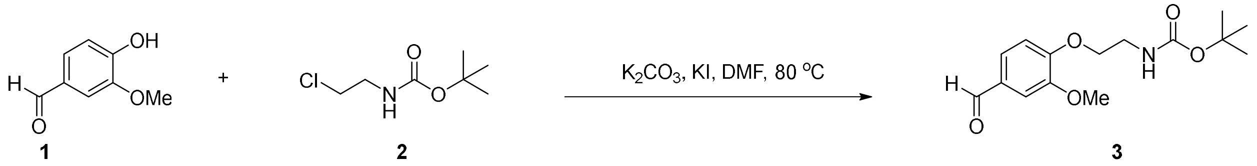

Compound **1** (10.0 g, 65.7 mmol), potassium carbonate (13.6 g, 98.6 mmol) and potassium iodide (2.2 g, 13.1 mmol) were suspended in DMF (150 ml), and then slowly added compound **2** (15.4 g, 85.5 mmol) in DMF (50 mL). The reaction mixture was stirred at 80°C for 20 h. At this point, TLC (silica, 1:3 EtOAc–hexanes) showed that the reaction was complete. The reaction mixture was quenched with cold water (300 mL), and the product was extracted with ethyl acetate (300 mL) three times, and then the organic layer was washed with 1M NaCl (300 mL) three times. The ethyl acetate layer was evaporated under reduced pressure to obtain compound **3** (16.3 g, yield 84%). $^1$H NMR (600 MHz, $CDCl_3$) δ 9.88 (s, 1H), 7.46 (dd, $J_1$ = 1.8 Hz, $J_2$ = 8.4 Hz, 1H), 7.44 (s, 1H), 7.01 (d, $J$ = 8.4 Hz, 1H), 5.12 (s, 1H), 4.18 (t, $J$ = 4.8 Hz, 2H), 3.95 (s, 3H), 3.62 (t, $J$ = 4.2, 2H), 1.49 (s, 9H); $^{13}$C NMR (150 MHz, $CDCl_3$) δ 190.9, 154.1, 149.9, 130.1, 126.6, 111.9, 109.3, 70.1, 61.7, 55.9, 25.8, 18.3.

### Synthesis of compound 5

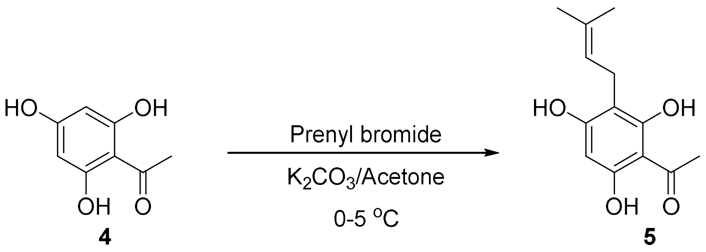


Compound **4** (10.0 g, 59.5 mmol) was dissolved in acetone (200 mL) followed by addition of $K_2CO_3$ (8.2 g, 59.5 mmol) and prenyl bromide (6.2 g, 41.6 mmol) at 0-5 °C. The reaction mixture continued for 2 hours at the same temperature. At this point, TLC (silica, 1:2 EtOAc–hexanes) showed that the reaction was complete. The reaction mixture was filtered, and the solvent was evaporated under reduced pressure. The residue was purified by column chromatography eluted with ethyl acetate/hexanes gradient to afford the desired product **5** (4.4 g, yield 31%) as a white solid. $^1$H NMR (600 MHz, Methanol-*d4*) δ 5.85 (s, 1H), 5.13 (t, *J* = 6.6 Hz, 1H), 3.06 (d, *J* = 7.2 Hz, 2H), 2.69 (s, 3H), 1.69 (s, 3H), 1.59 (s, 3H); $^{13}$C NMR (150 MHz, Methanol-*d4*) δ 204.6, 164.7, 163.9, 161.8, 131.1, 124.5, 107.9, 105.5, 94.8, 32.8, 25.9, 22.1, 17.8.

**Synthesis of compound 6**

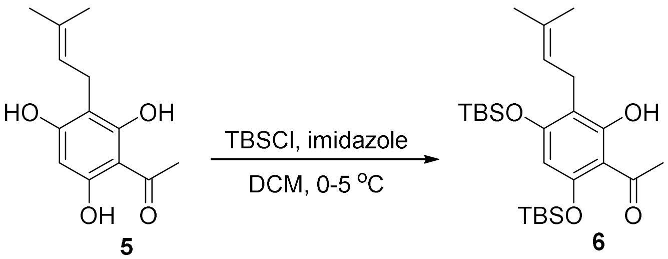


Compound **5** (5.0 g, 21.2 mmol) was dissolved in DCM (50 mL) and imidazole (5.8 g, 63.5 mmol) at 0-5 °C. *Tert*-butyldimethylsilyl chloride (6.4 g, 42.3 mmol) was slowly added, and then the reaction temperature was slowly raised to room temperature and continued for 3h. After consumption of the starting material (by TLC), the reaction mass was quenched with ice cold water (200 mL), and the product was extracted with dichloromethane (60 mL) three times, and then the organic layer was washed with 1M $NaHCO_3$ (60 mL) three times. The ethyl acetate layer was evaporated under reduced pressure, and then the residue was purified by column chromatography eluted with ethyl acetate/hexanes gradient to afford the desired product **6** (8.7 g, yield 88%) as a yellow oil. $^1$H NMR (600 MHz, $CDCl_3$) δ 13.76 (s, 1H), 5.85 (s, 1H), 5.14 (t, *J* = 6.6 Hz, 1H), 3.25 (d, *J* = 6.6 Hz, 2H), 2.62 (s, 3H), 1.75 (s, 3H), 1.66 (s, 3H), 1.01 (s, 18H), 0.34 (s, 6H), 0.27 (s,

6H); $^{13}$C NMR (150 MHz, $CDCl_3$) δ 203.4, 164.0, 159.7, 157.1, 131.2, 123.0, 113.3, 108.7, 100.9, 32.9, 26.0, 25.7, 25.6, 21.9, 18.8, 18.3, 17.9, -3.5, -4.0.

**Synthesis of compound 7**

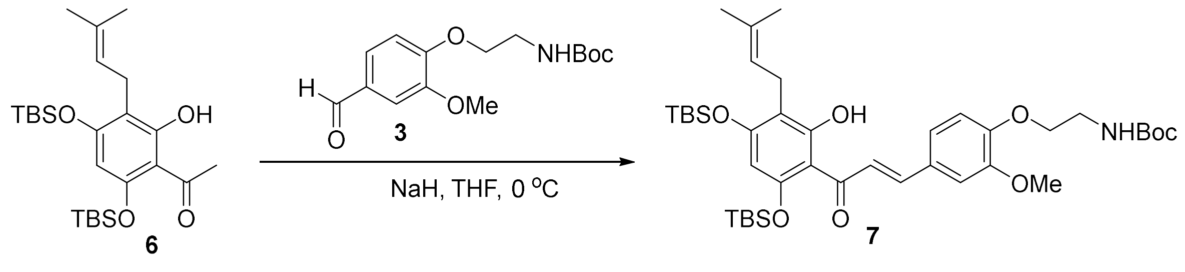


Compound **6** (2.0 g, 4.3 mmol) was dissolved in THF (60 mL) at 0-5 °C, and then slowly added NaH (0.12 g, 5.2 mmol) and stirred for 10-15 min, followed by addition of compound **3** (1.4 g , 4.7 mmol) in THF (6 mL) dropwise at 5-10 °C. Progress of the reaction was monitored by TLC. After completion of starting material (by TLC), the reaction mass was quenched with Ammonium chloride (100 mL) at 0 °C and the product was extracted with ethyl acetate (100 mL) three times. The organics layer was evaporated, and the residue was purified by column chromatography eluted with ethyl acetate/hexanes gradient to afford the desired product **7** (2.3 g, yield 79%) as a yellow oil. $^{1}$H NMR (600 MHz, $CDCl_3$) δ 13.21 (s, 1H), 7.73 (d, $J$ = 14.4 Hz, 1H), 7.58 (d, $J$ = 14.4 Hz, 1H), 7.15 (dd, $J_1$ = 1.8 Hz, $J_2$ = 8.4 Hz, 1H), 7.13 (d, $J$ = 1.8 Hz, 1H), 6.88 (d, $J$ = 8.4 Hz, 1H), 5.89 (s, 1H), 5.15 (m, 2H),4.11 (t, $J$ = 4.8 Hz, 2H), 3.89 (s, 3H), 3.57 (d, $J$ = 6.6 Hz, 2H), 3.27 (d, $J$ = 6.6 Hz, 2H), 1.75 (s, 3H), 1.67 (s, 3H), 1.46 (s, 9H), 0.92 (s, 18H), 0.34 (s, 6H), 0.27 (s, 6H); $^{13}$C NMR (150 MHz, $CDCl_3$) δ 193.3, 163.4, 159.6, 156.0, 149.6, 142.0, 131.3, 129.1, 126.0, 123.2, 123.1, 113.8, 110.3, 102.3, 55.8, 28.4, 25.9, 25.7, 25.6, 22.1, 18.6, 17.9, -0.36, -0.39.

**Synthesis of compound 8**

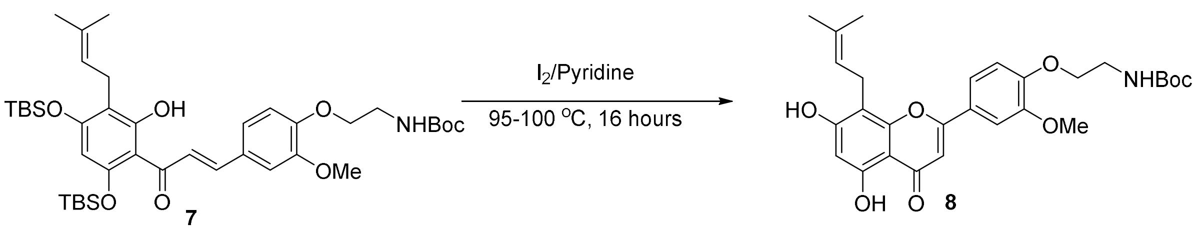


Compound **7** (1.0 g, 1.35 mmol) was suspended in 100 mL of pyridine followed by addition of Iodine (0.38 g, 1.48 mmol) and raised the temperature to 95-100 °C for 16 h. Progress of the reaction was monitored by TLC. After completion of starting material (by TLC), the reaction mass was quenched with water (30 mL) and reaction mass pH was adjusted to 3-4 by using citric acid. The crude product was washed with 10% $Na_2S_2O_3$ solution, extracted with EtOAc (100 mL) three times and washed with the organic layer with water (100 mL) three times. Organic layer was dried over sodium sulphate and evaporated the solvent under reduced pressure to get crude product as yellow colored solid. The organics layer was evaporated, and the residue was purified by column chromatography eluted with ethyl acetate/hexanes gradient to afford the desired product **8** (0.21 g, yield 30%) as yellow solid. $^1$H NMR (600 MHz, $CDCl_3$) δ 7.41 (d, $J$ = 6.8 Hz, 1H), 7.30 (s, 1H), 6.89 (d, $J$ = 9.0 Hz, 1H), 6.54 (s, 1H), 6.34 (s, 1H), 5.29 (t, $J$ = 6.6 Hz, 1H), 5.21 (s, 1H), 4.11 (t, $J$ = 5.4 Hz, 2H), 3.91 (s, 3H), 3.53 (d, $J$ = 6.0 Hz, 2H), 1.82 (s, 3H), 1.71 (s, 3H), 1.47 (s, 9H); $^{13}$C NMR (150 MHz, DMSO-*d6*) δ 182.6, 163.6, 162.1, 159.5, 156.1, 155.0, 151.7, 149.6, 131.6, 123.9, 122.9, 120.3, 113.5, 110.3, 106.6, 104.2, 104.1, 98.9, 78.3, 67.6, 56.2, 28.7, 25.8, 21.8, 18.3.

**Synthesis of compound 9**

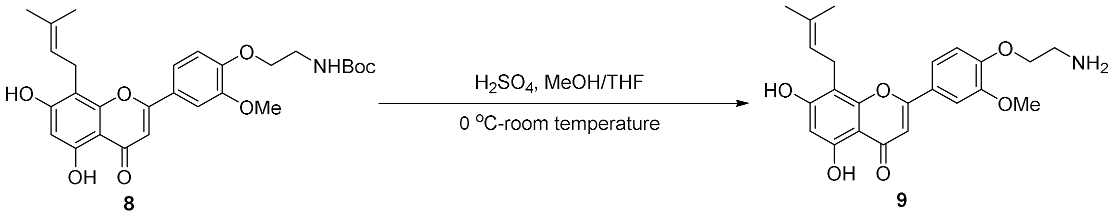


Compound **8** (300 mg, 0.72 mmol) was suspended in MeOH/THF (1/5 ratio, 1.8 mL) at 0-5 °C, and then slowly added $H_2SO_4$ (0.3 mL). The resultant mixture was slowly raised to room temperature and continued for 3h. Completion of reaction is monitored by TLC (10% MeOH:DCM: and 3-4 drops of aq. ammonia). Quenched the reaction by slow addition of ice cold water and stirred for 30 min. Filtered the solid and washed with EtOAc, dried under vacuum at 40 °C to get yellow crude product. The crude product was purified by column chromatography eluted with acetone/hexanes (1% $Et_3N$) gradient to afford the desired product **9** (50 mg, yield 21%) as yellow solid. $^1$H NMR (600 MHz, DMSO-*d6*) δ 7.62 (dd, $J_1$ = 1.8 Hz, $J_2$ = 8.4 Hz, 1H), 7.56 (d, $J$ = 1.8 Hz, 1H), 7.16 (d, $J$ = 9.0 Hz, 1H), 6.93 (s, 1H), 6.26 (s, 1H), 5.23 (t, $J$ = 6.6 Hz, 1H), 4.05 (t, $J$ = 6.0 Hz, 2H), 3.87 (s, 3H), 2.95 (t, $J$ = 6.0 Hz, 2H), 1.75 (s, 3H), 1.63 (s, 3H); ); $^{13}$C NMR (150 MHz, DMSO-*d6*) δ 182.6, 163.6, 162.0, 159.5, 156.1, 155.0, 152.0, 149.5, 131.6, 123.6, 122.9, 120.3, 113.1, 109.9, 106.6, 104.2, 104.0, 98.9, 70.8, 59.8, 56.1, 25.8, 21.8, 18.3.

**Final step to Caflanone-PEG(3400)-DSPE conjugate**

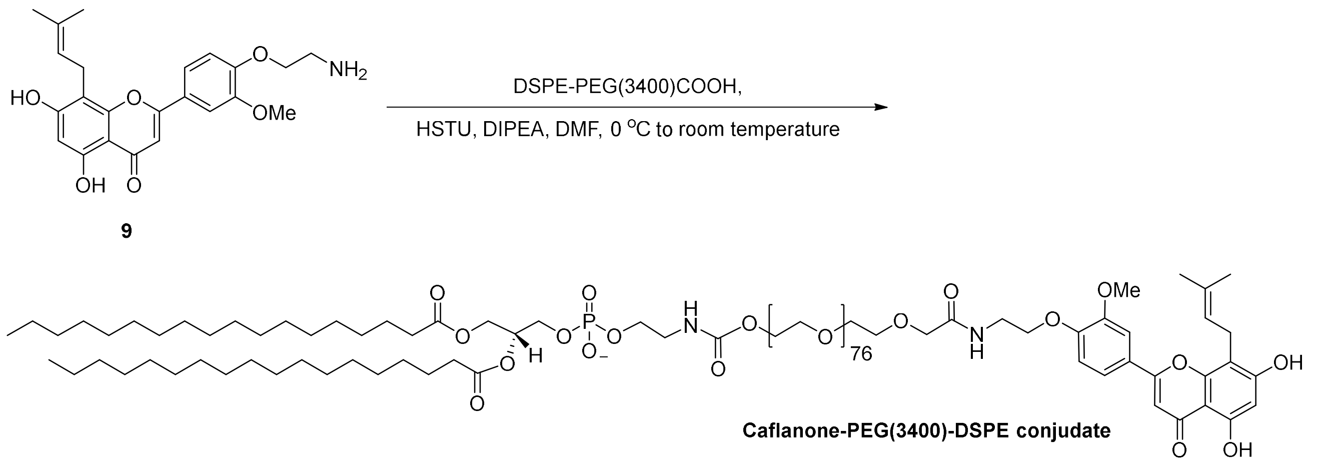


To a solution of DSPE-PEG(3400)-COOH (500 mg, 0.12 mmol) and HLNT-06 (193 mg, 0.40 mmol) in dry DMF (8 mL) was added HSTU (160.0 mg, 0.40 mmol). The reaction mixture was stirred at room temperature for two days and then concentrated under reduced pressure. The residue was diluted in methanol/water mixture (1:1, 8 ml), loaded into a 2000 MWCO dialysis bag and dialyzed against MES buffer (50 mM, 2 × 5 liters) for 8 hours and then water (3 × 5 liters) for 2 days. The water was then removed by freeze drying to obtain the desired compound as a yellow solid (253 mg, 47%). $^1$H NMR (600 MHz, $CDCl_3$) δ 12.67 (s, 1H), 11.97 (s, 1H), 7.47 (d, *J* = 8.4 Hz, 1H), 7.38 (s, 1H), 6.97 (d, *J* = 8.4 Hz, 1H), 6.54 (s, 1H), 6.46 (s, 1H), 5.29-5.27 (m, 1H), 5.22-5.19 (m, 1 H), 4.55-4.53 (m, 2 H), 4.35 (dd, $J_1$ = 3.6 Hz, $J_2$ = 12.0 Hz, 1H), 4.33-4.31 (m, 2 H), 3.91 (s, 3H), 3.76-3.50 (m, ~360 H), 3.06-3.05 (m, 4 H), 2.26 (t, *J* = 7.8 Hz, 6H), 1.78 (s, 3H), 1.67 (s, 3H), 1.59-1.55 (m, 4 H), 1.32-1.23 (m, 60 H), 0.86 (t, *J* = 7.2 Hz, 6H). HRMS (MALDI) *calcd* for $C_{221}H_{415}N_2O_{94}P$ $[M+NH_4]^+$ 4652.640, found, 4652.633.

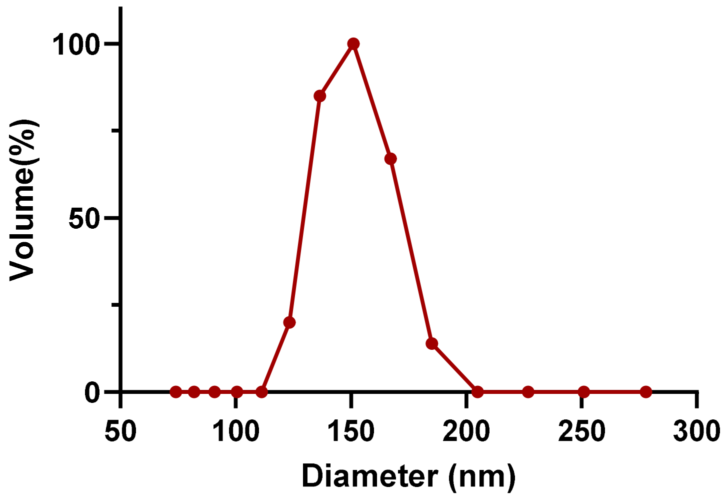


**Fig. S1**. The particle size of **NCDx** was measured using Dynamic Light Scattering (DLS). The results showed an average hydrodynamic diameter of **148.95 ± 26.63 nm**.

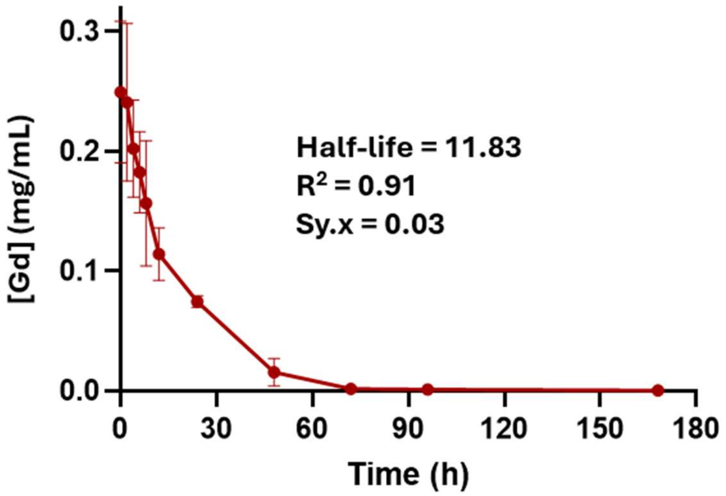


**Fig. S2**. The systemic circulation half-life of **NCDx** was evaluated in 8-10 weeks old C57BL/6 mice. The agent was administered via tail vein injection (dose 0.2 mmol Gd(III)/kg) and blood samples were collected from three mice at different time points over a period of 1 week and analyzed for gadolinium (Gd) content. The blood Gd concentrations were quantified using inductively coupled plasma mass spectrometry **(ICP-MS)**. The elimination half-life (t½) was determined to be 11.83 h ($R^2$ = 0.91, Sy.x = 0.03) using GraphPad Prism.

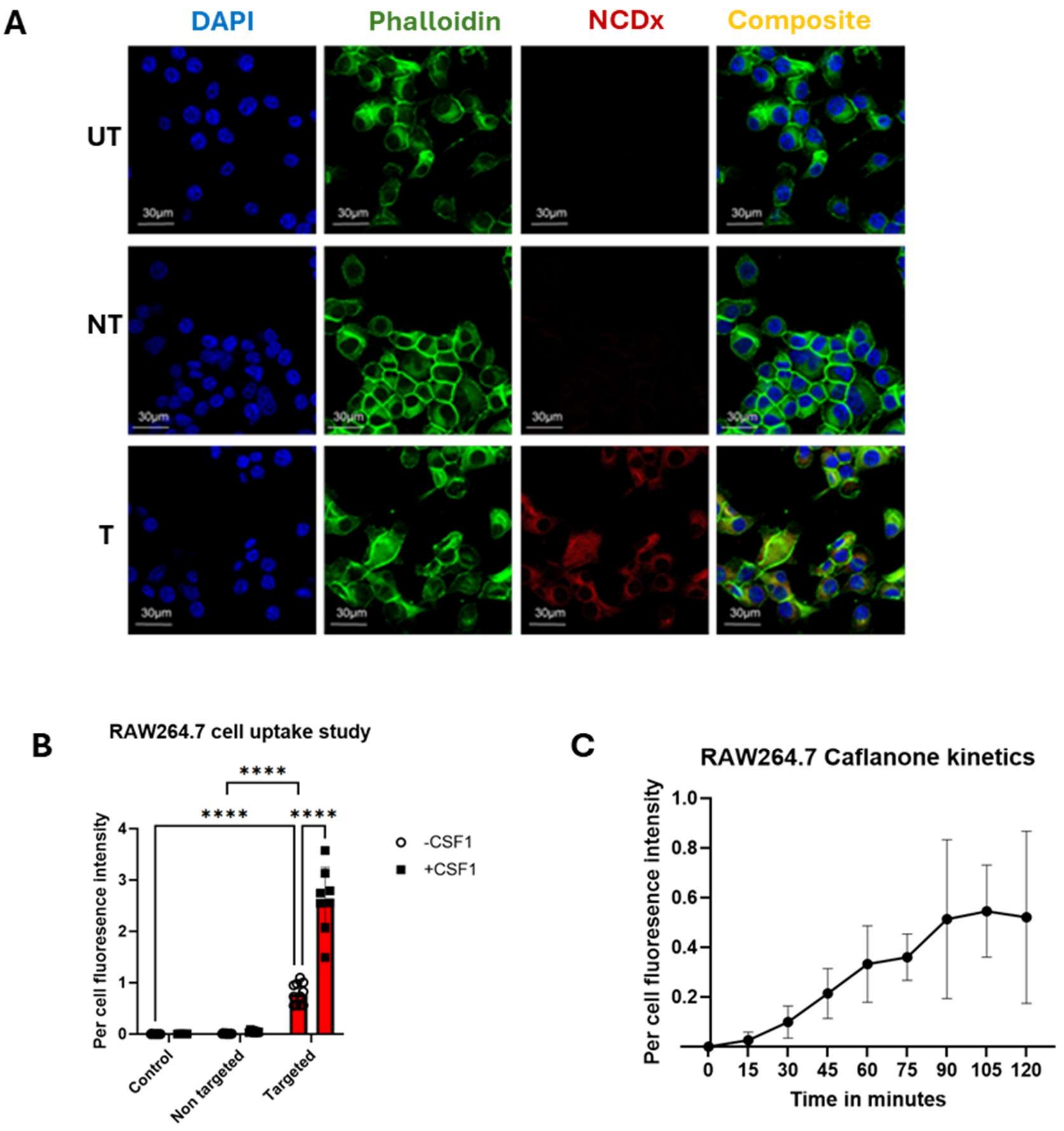


**Fig. S3**. Raw 264.7 cell line actively internalize **NCDx**. **(A**) Raw 264.7 cells show avid internalization of **NCDx** within 2 hours of exposure; (**B**) Activation of Raw 264.7 cells with CSF1 results in increased uptake of **NCDx**; (**C**) A plot of signal intensity per cell against exposure time demonstrating that **NCDx** internalization exhibits saturation kinetics. Scale bar = 30 µm.

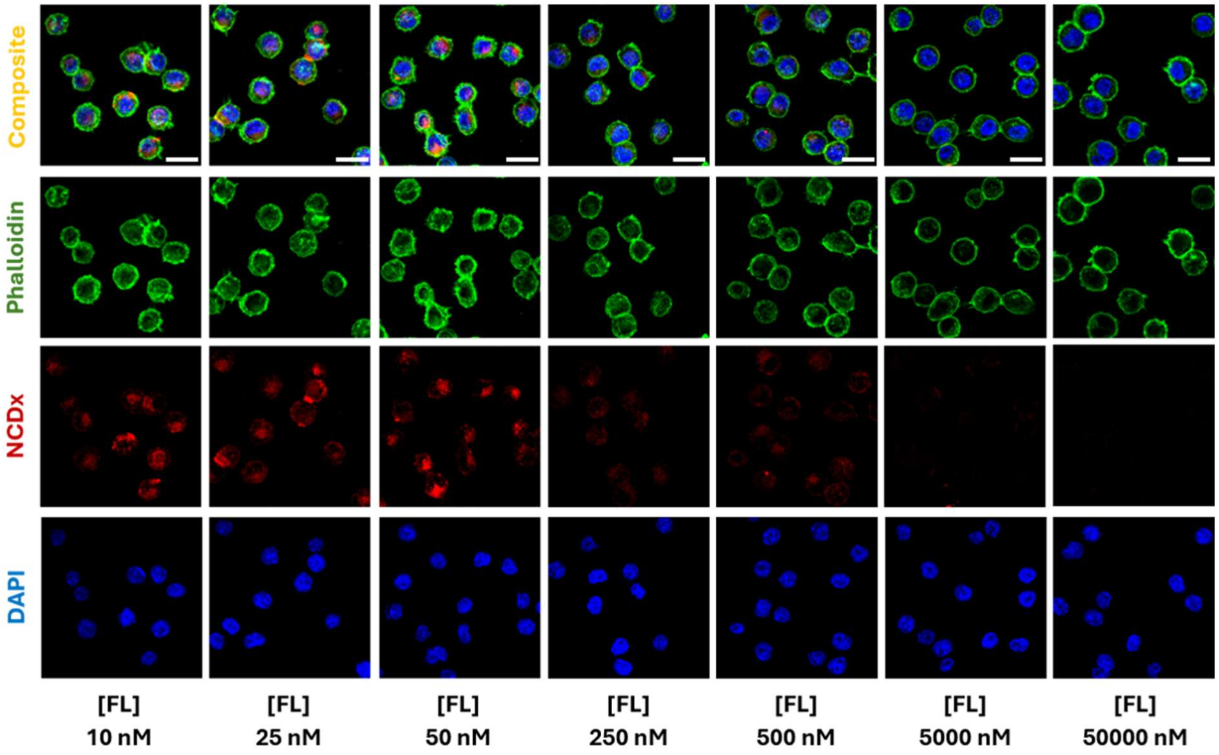


**Fig. S4.** Sim A9 cell blocking study with different concentrations of free ligand (Caflanone). Scale bar = 20 μm.

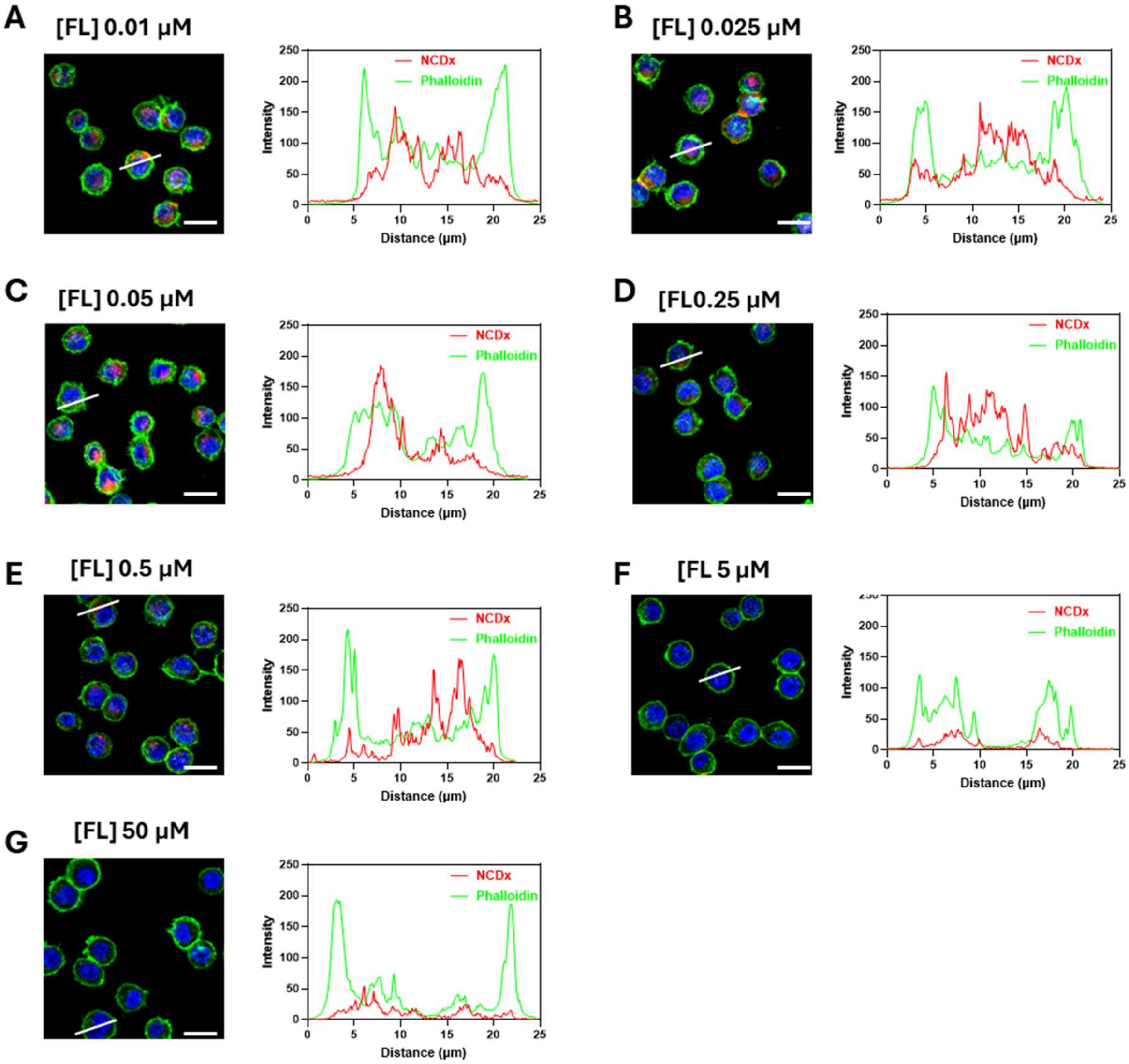


**Fig. S5.** Sim A9 cell blocking study with different concentration of free ligand (Caflanone). **A** to **G** is Intensity curve along the indicative profile line in confocal images. Scale bar = 20 µm.

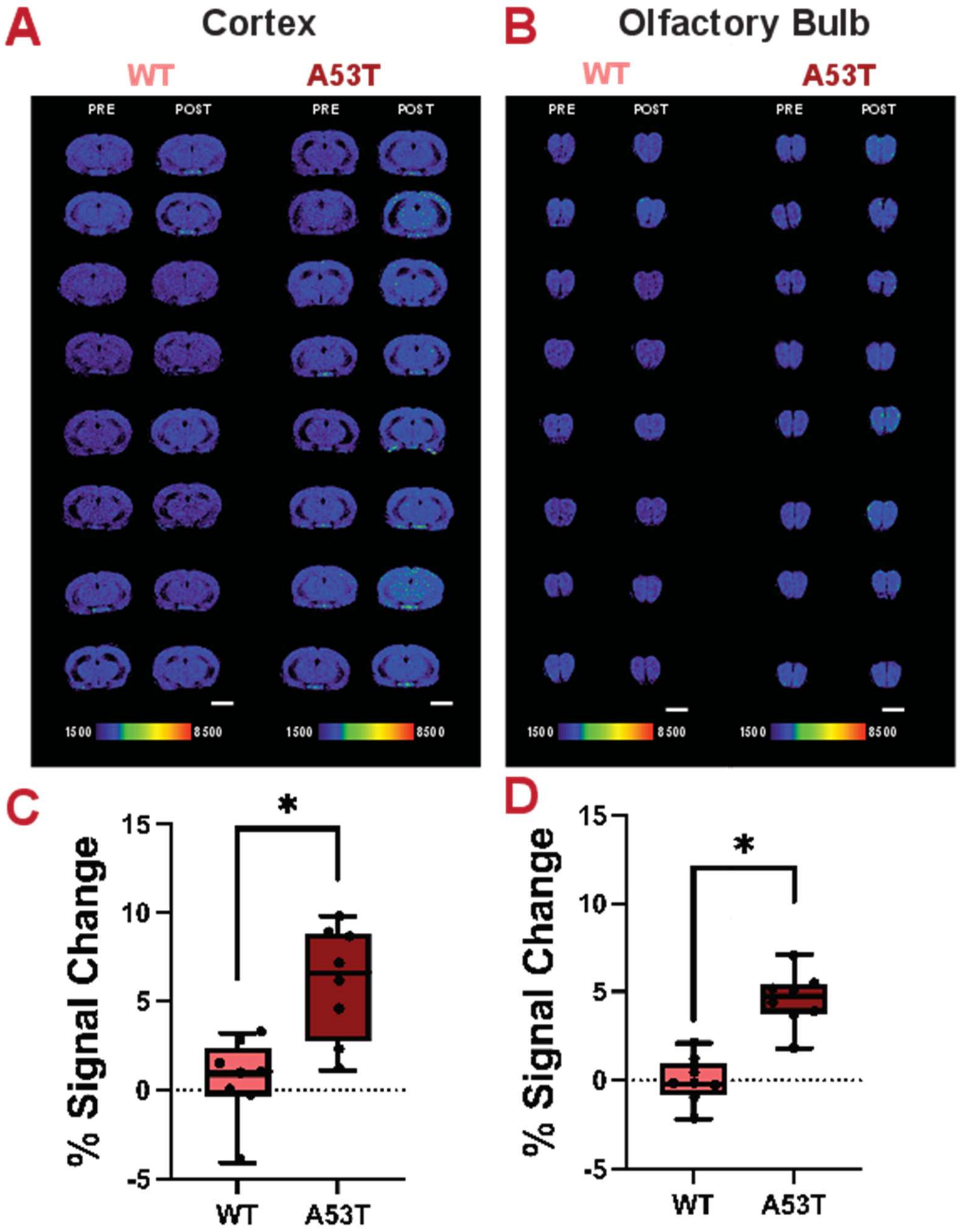


**Fig. S6**: **In *vivo* contrast-enhanced MRI of all mice injected with NCDx.** (**A**) Axial 3.6 mm thick slabs cortices (Bregma 0 mm) and (**B**) olfactory bulbs (Bregma 4-5 mm) of pre- and post-contrast images of TG mice using a jet colormap to illustrate particle label intensity show apparent increase in signal intensity in POST scans compared to PRE. All images were analyzed on the same MR signal intensity scale (1500-4000 a.u.); (**C**) Box and whisker plot quantification of percent change in signal intensity demonstrate statistically signal increase in TG versus WT mice cortices; (**D**) Box and whisker plot quantification of percent change in signal intensity demonstrate statistically signal increase in TG versus WT mice olfactory olfactory bulbs. Wilcoxon rank sum test: $^*p<0.05$. Scale bar = 3 mm

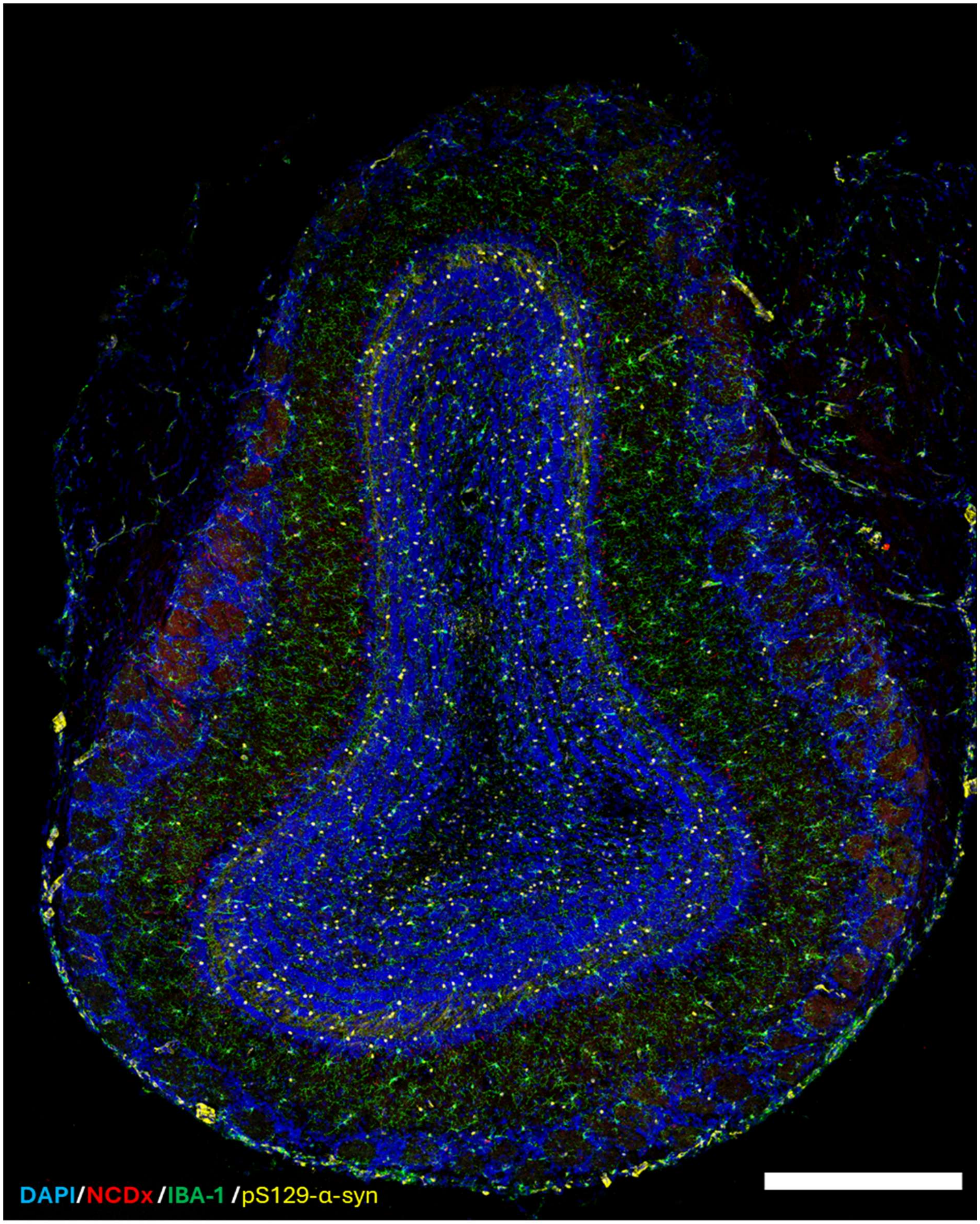


**Fig. S7**. The **DAPI/NCDx/IBA-1/pS129-α-syn** composite confocal image demonstrates that *in vivo* accumulation of **NCDx** in the olfactory bulb of **TG** mice is primarily mediated by microglial uptake. Scale bar = 300 μm

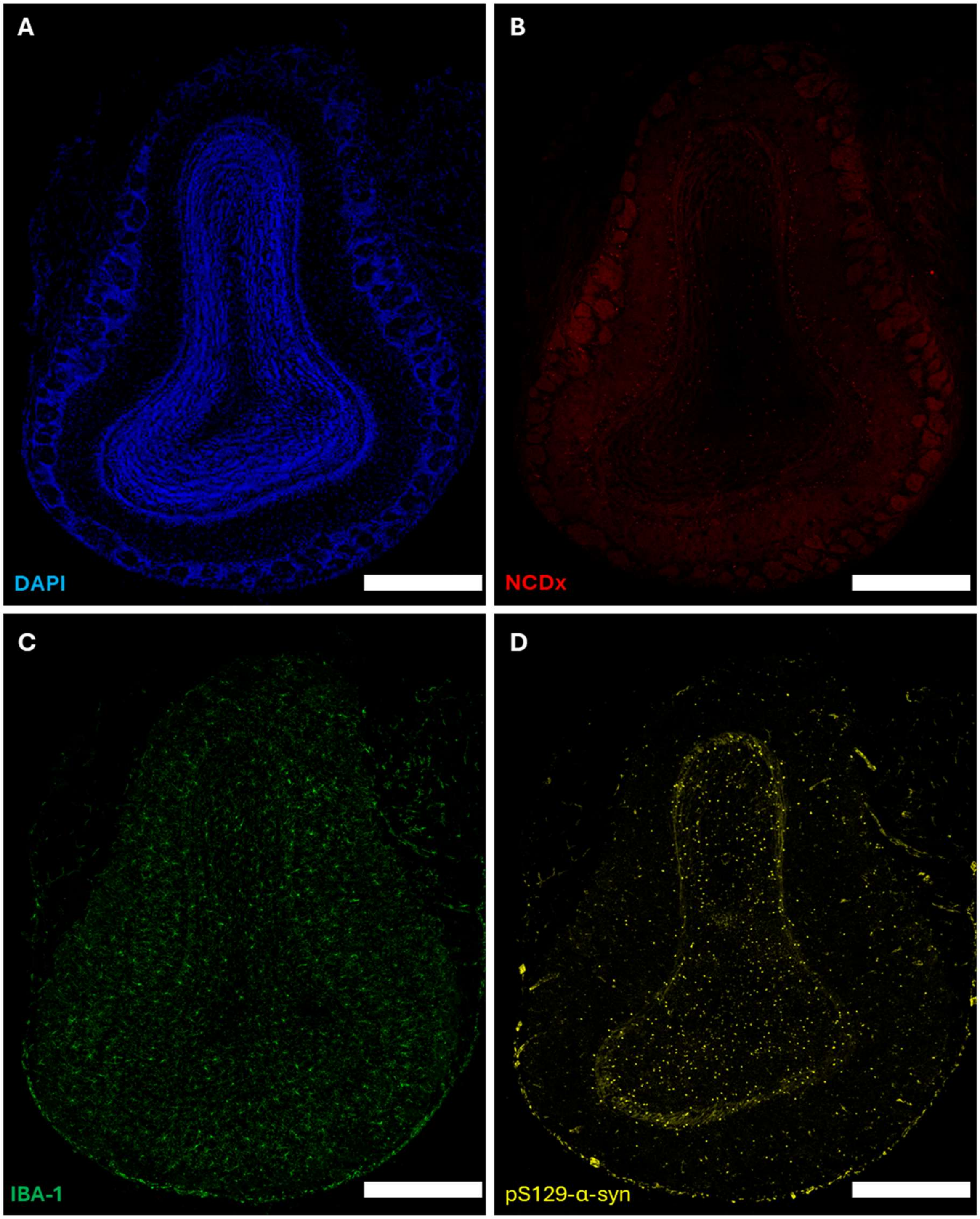


**Fig. S8**. Whole-section confocal image of the olfactory bulb from a transgenic (**TG**) mouse. (**A**) DAPI staining; (**B**) **NCDx** accumulation; (**C**) Anti-Iba1 staining; (**D**) anti-alpha-synuclein staining. Scale bar = 300 μm.

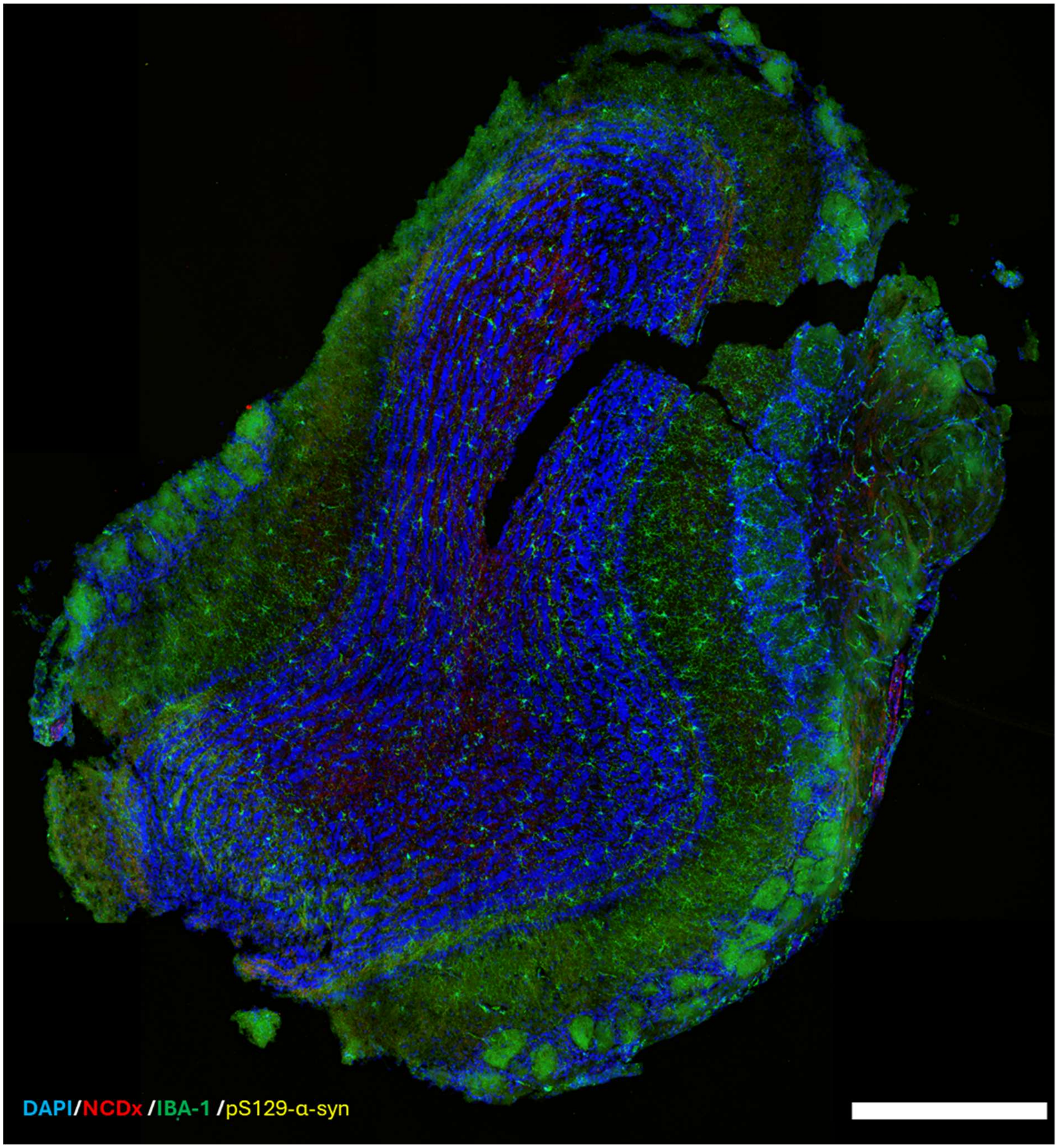


**Fig. S9**. The **DAPI/NCDx/IBA-1/pS129-α-syn** composite confocal image demonstrates that **NCDx** does not accumulate in microglial cells in the olfactory bulb of wild-type (WT) mice. Scale bar = 300 μm

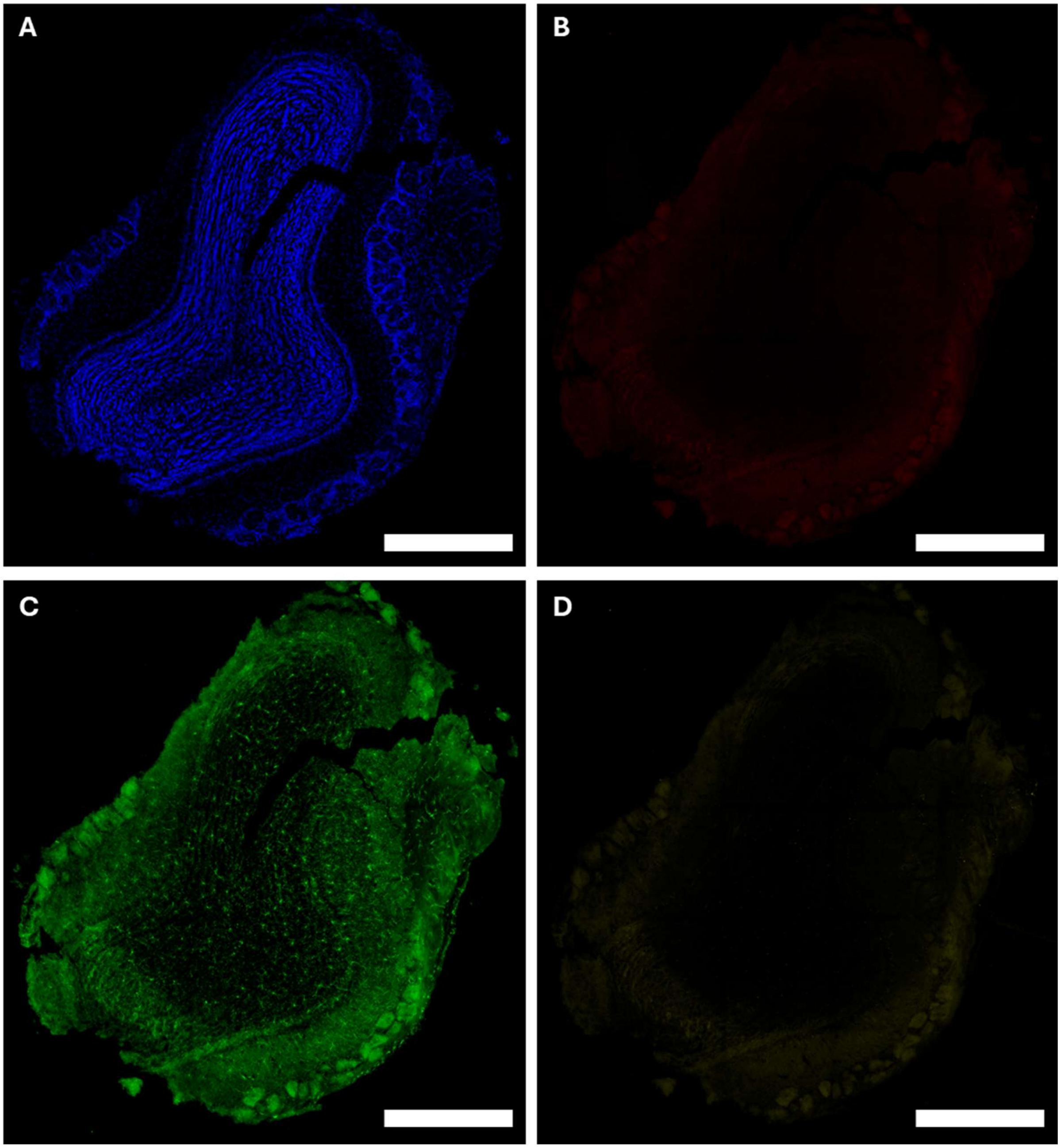


**Fig. S10**. Whole-section confocal image of the olfactory bulb from a wild-type (WT) mouse. (**A**) DAPI staining; (**B**) **NCDx** accumulation; (**C**) Anti-IBA-1 staining; (**D**) anti-alpha-synuclein staining. Scale bar = 300 µm.

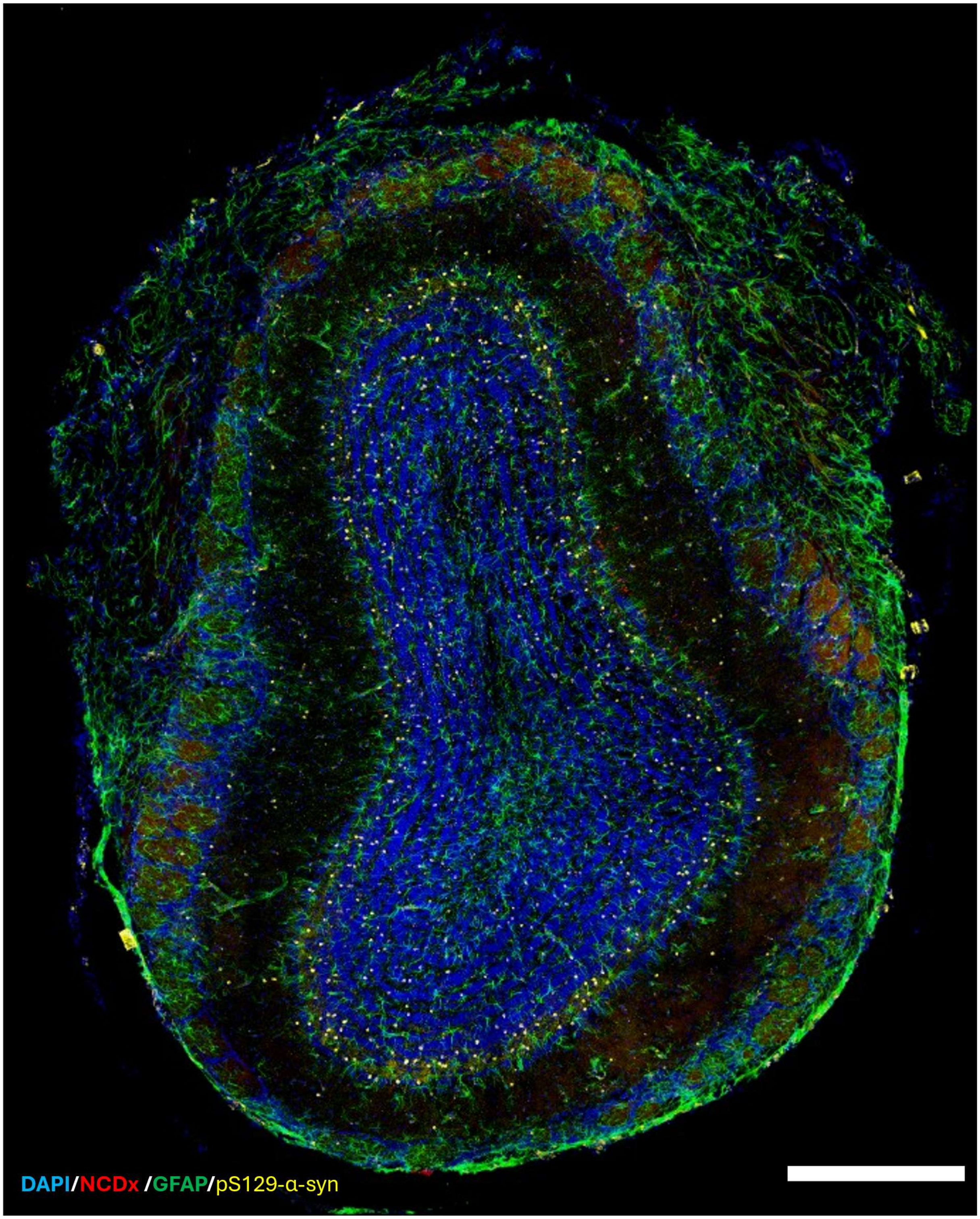


**Fig. S11.** The **DAPI/NCDx/GFAP/pS129-α-syn** composite confocal image demonstrates that **NCDx** does not accumulate in microglial cells in the olfactory bulb transgenic (TG) mouse. Scale bar = 300 μm

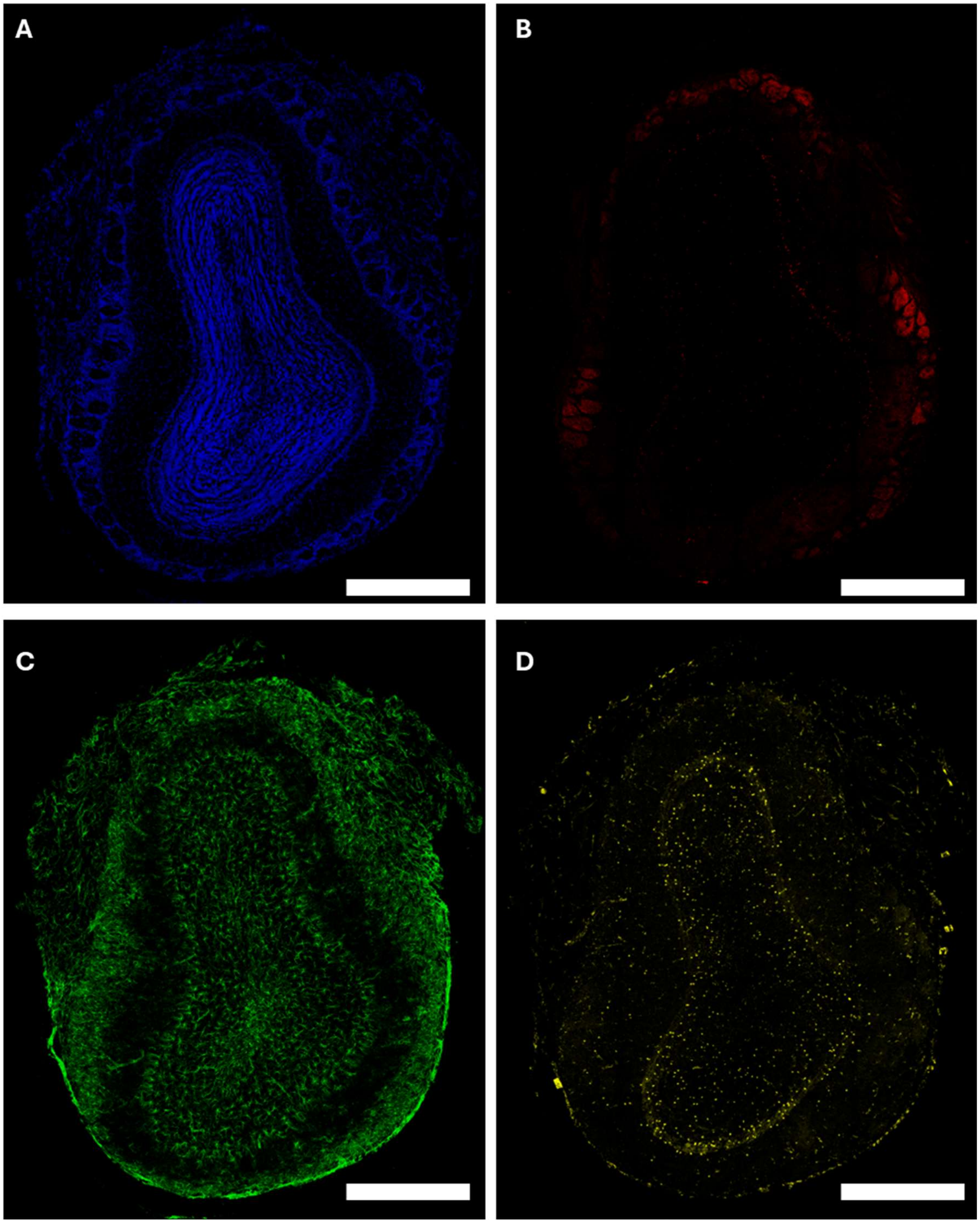


**Fig. S12.** Whole-section confocal image of the olfactory bulb from a transgenic (TG) mouse. (**A**) DAPI staining; (**B**) **NCDx** signal; (**C**) Anti-GFAP staining; (**D**) anti-alpha-synuclein staining. Scale bar = 300 µm.

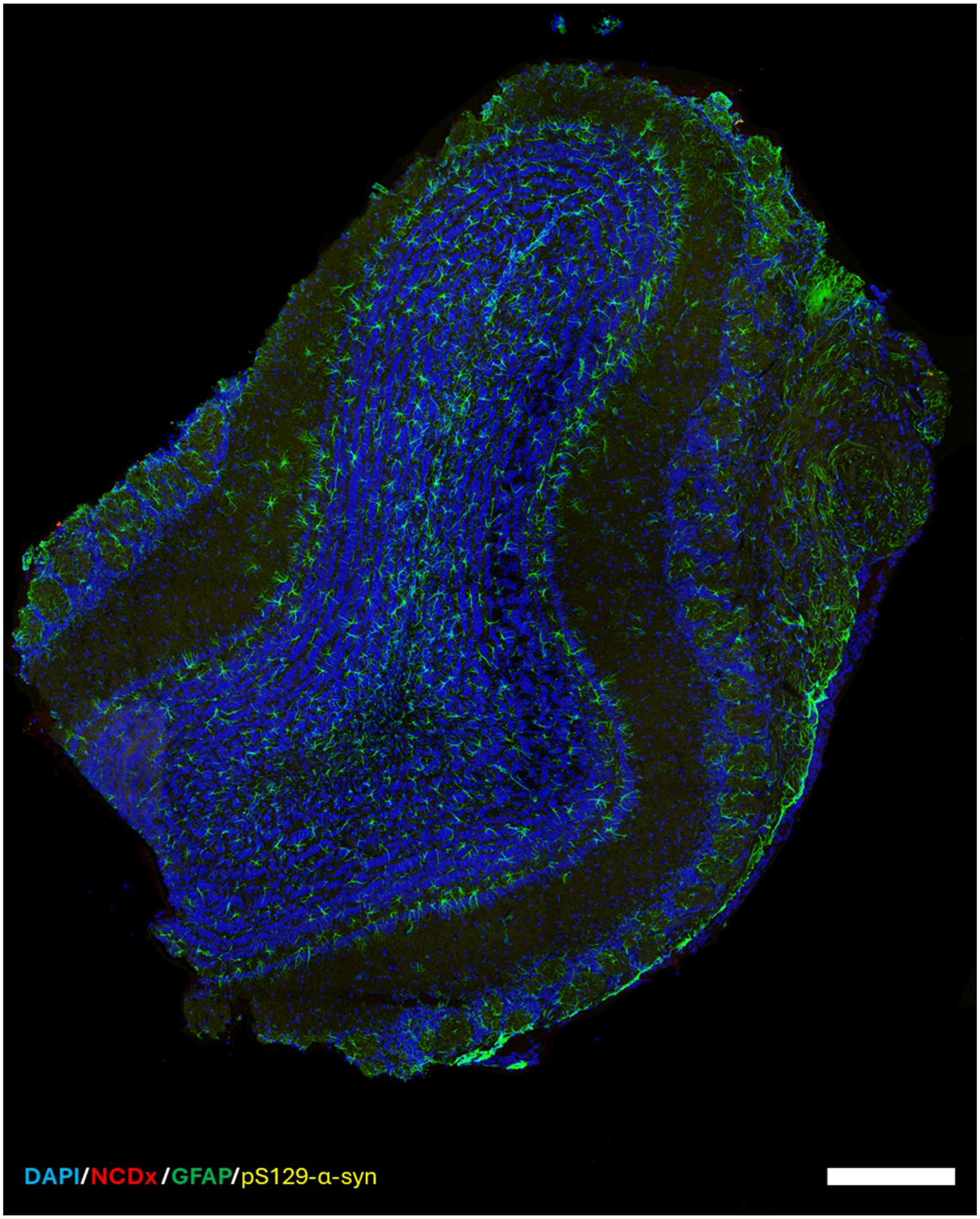


**Fig. S13**. The **DAPI/NCDx/GFAP/pS129-α-syn** composite confocal image demonstrates that **NCDx** does not accumulate in microglial cells in the olfactory bulb of wild-type (WT) mice. Scale bar = 300 μm

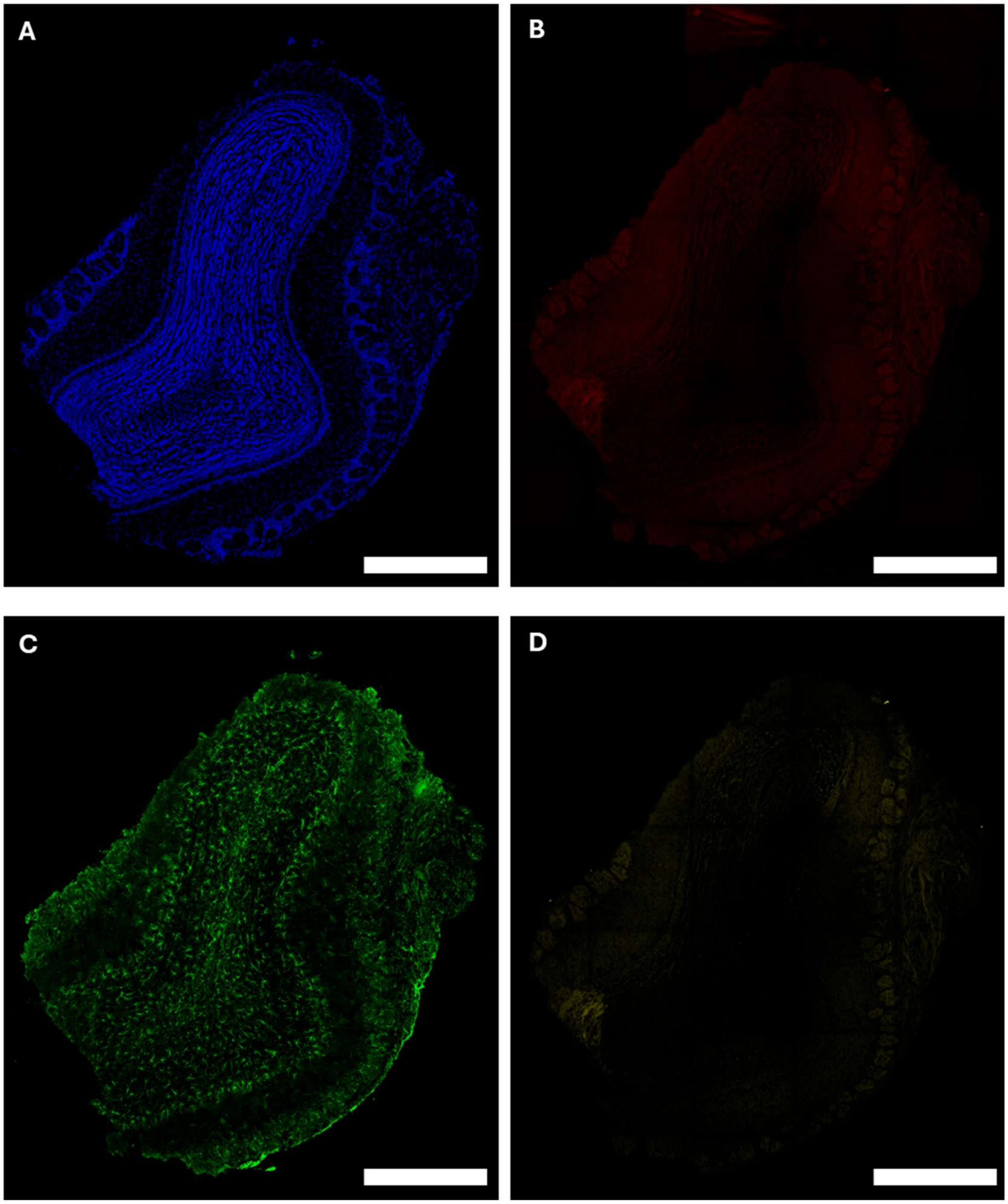


**Fig. S14**. Whole-section confocal image of the olfactory bulb from a wild-type (WT) mouse. (**A**) DPAI staining; (**B**) **NCDx** signal; (**C**) Anti-GFAP staining; (**D**) anti-alpha-synuclein staining. Scale bar = 300 μm.

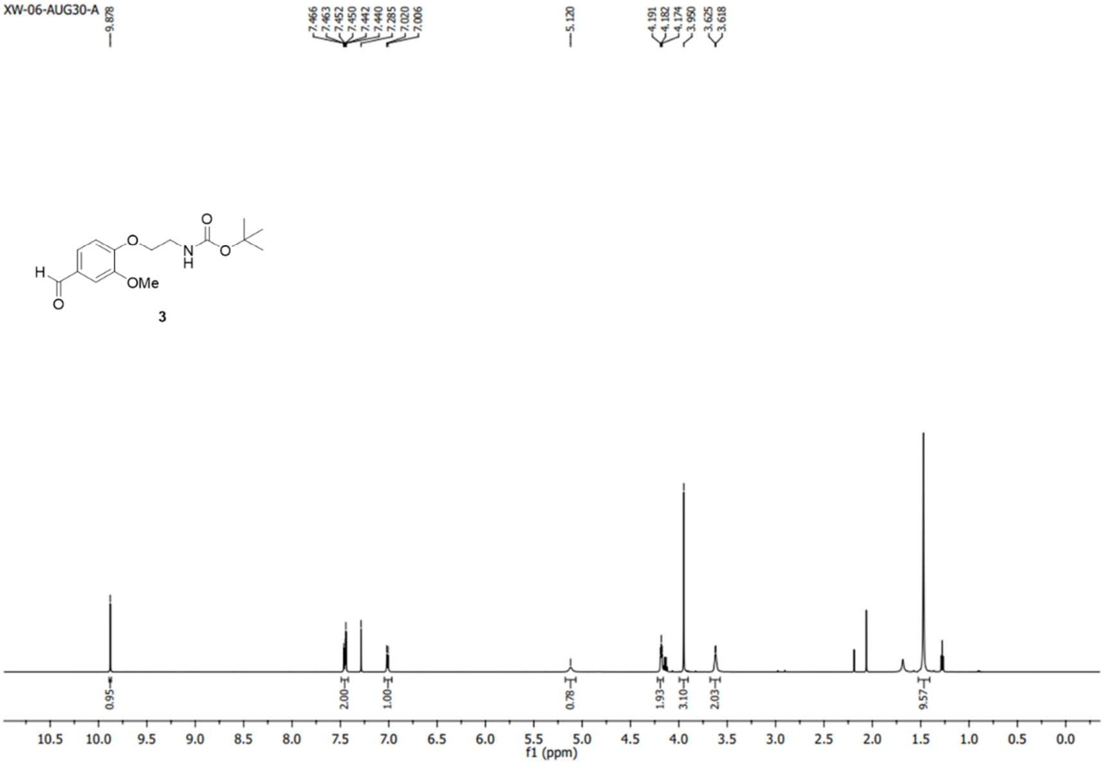


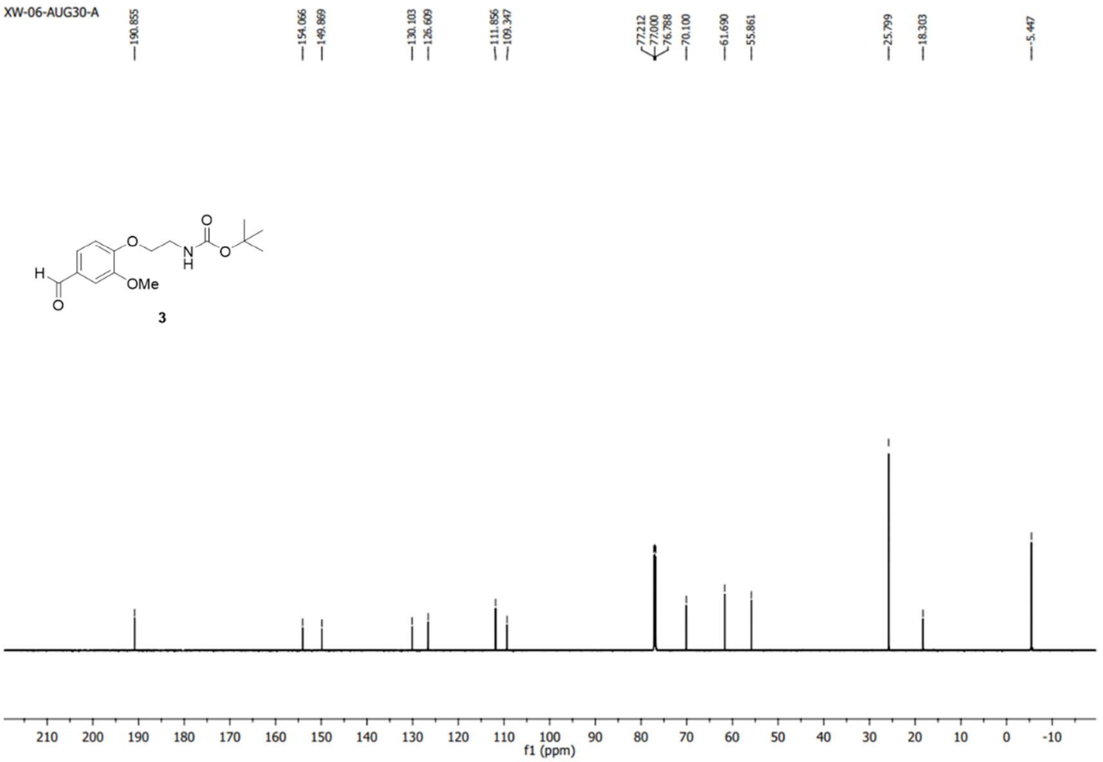


**Fig. S15**. $^{1}H$ and $^{13}C$ NMR Spectrum of Compound **3**

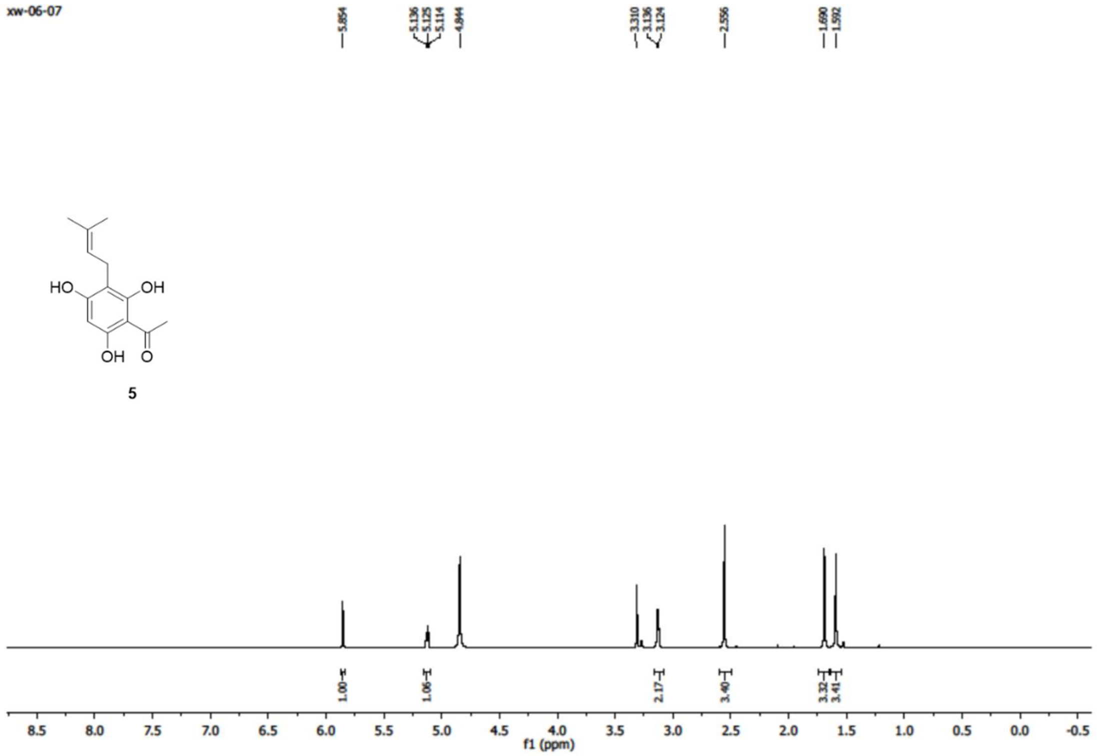


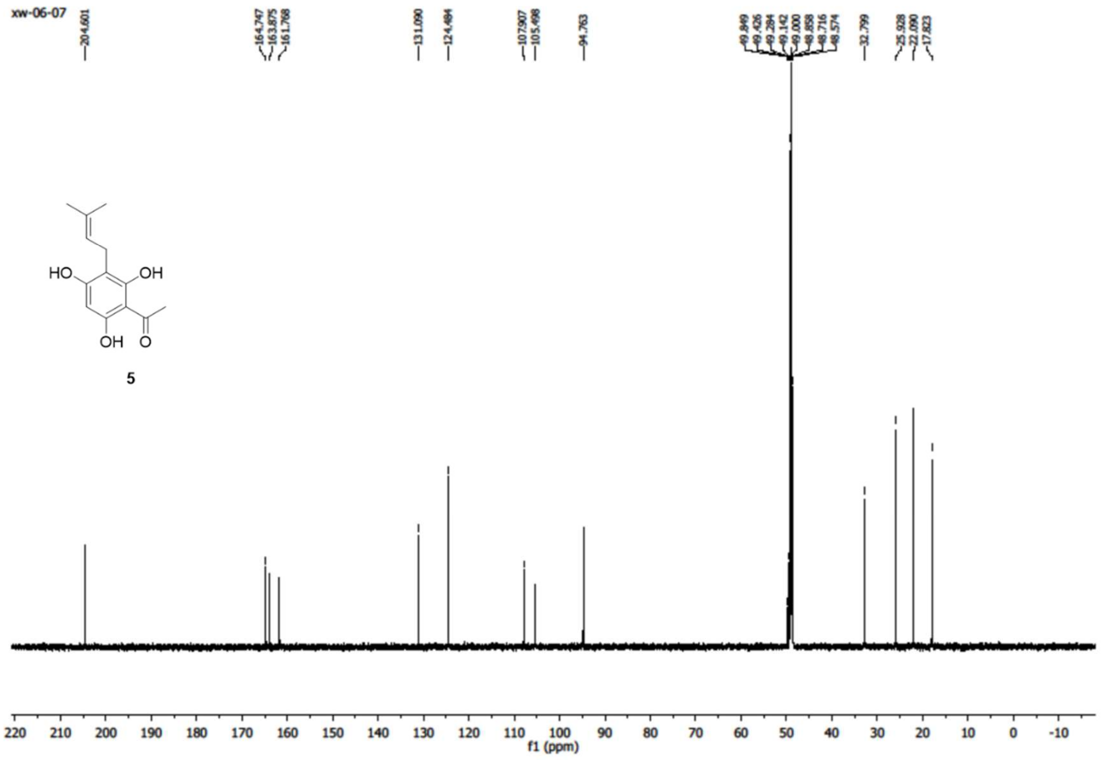


**Fig. S16**. [1]H and [13]C NMR Spectrum of Compound **5**

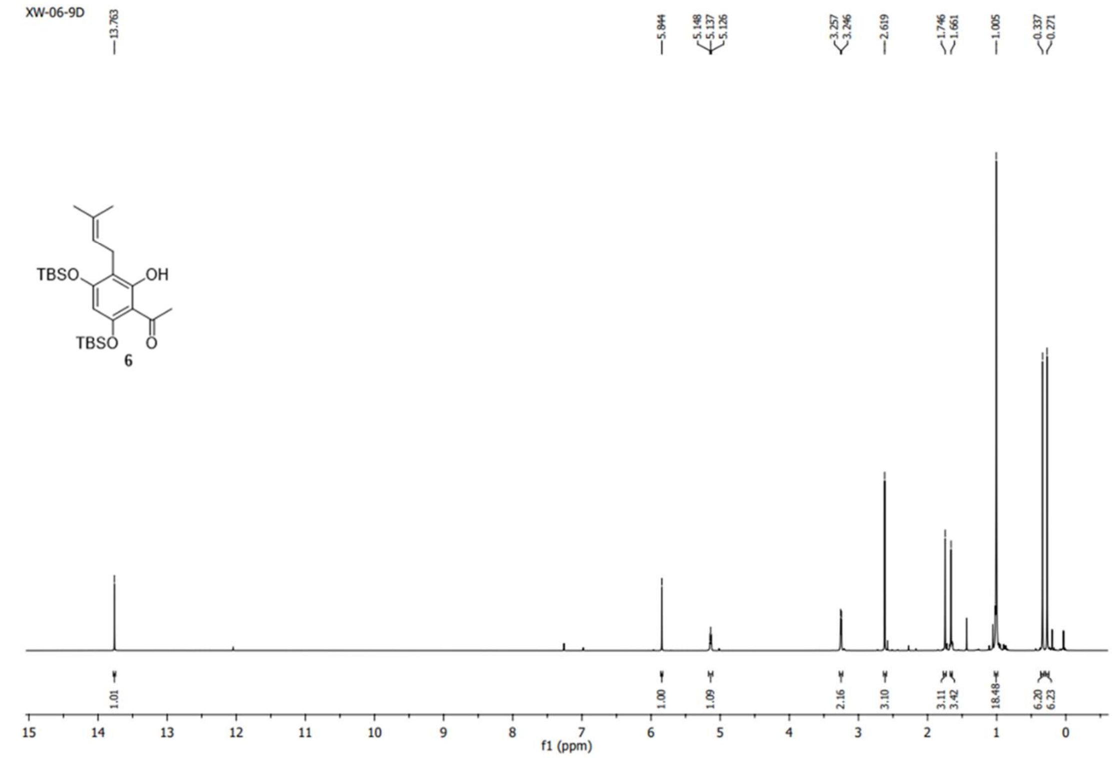


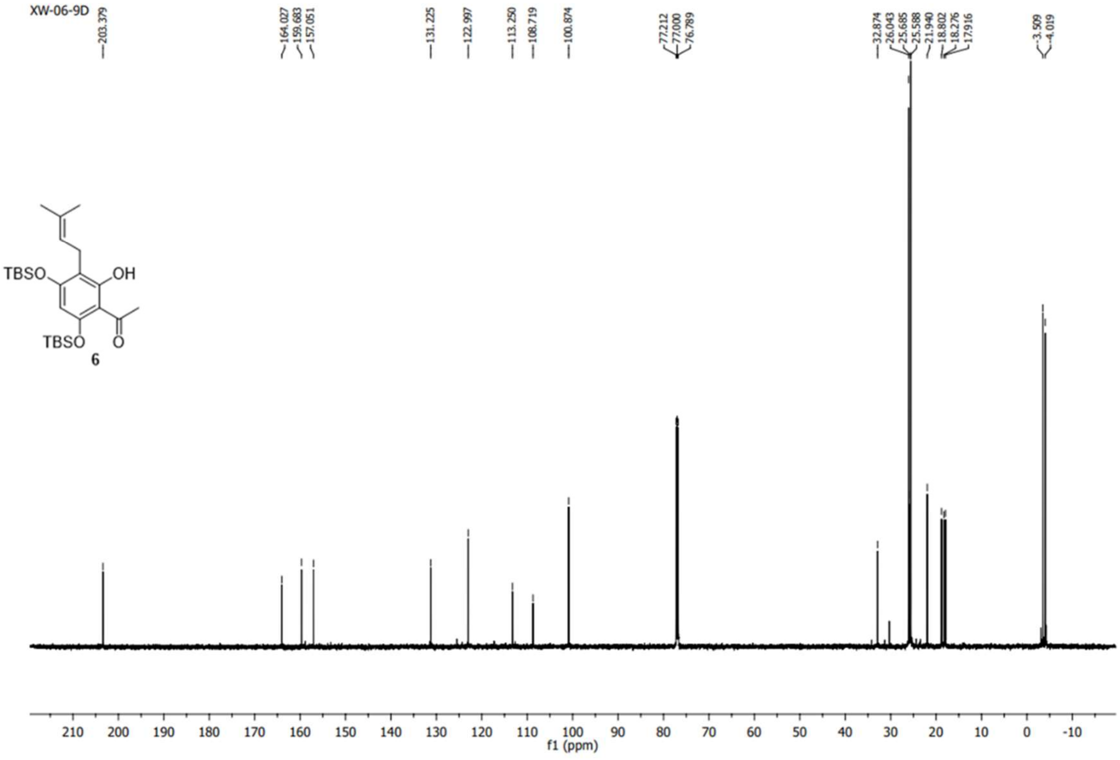


**Fig. S17**. $^{1}H$ and $^{13}C$ NMR Spectrum of Compound **6**

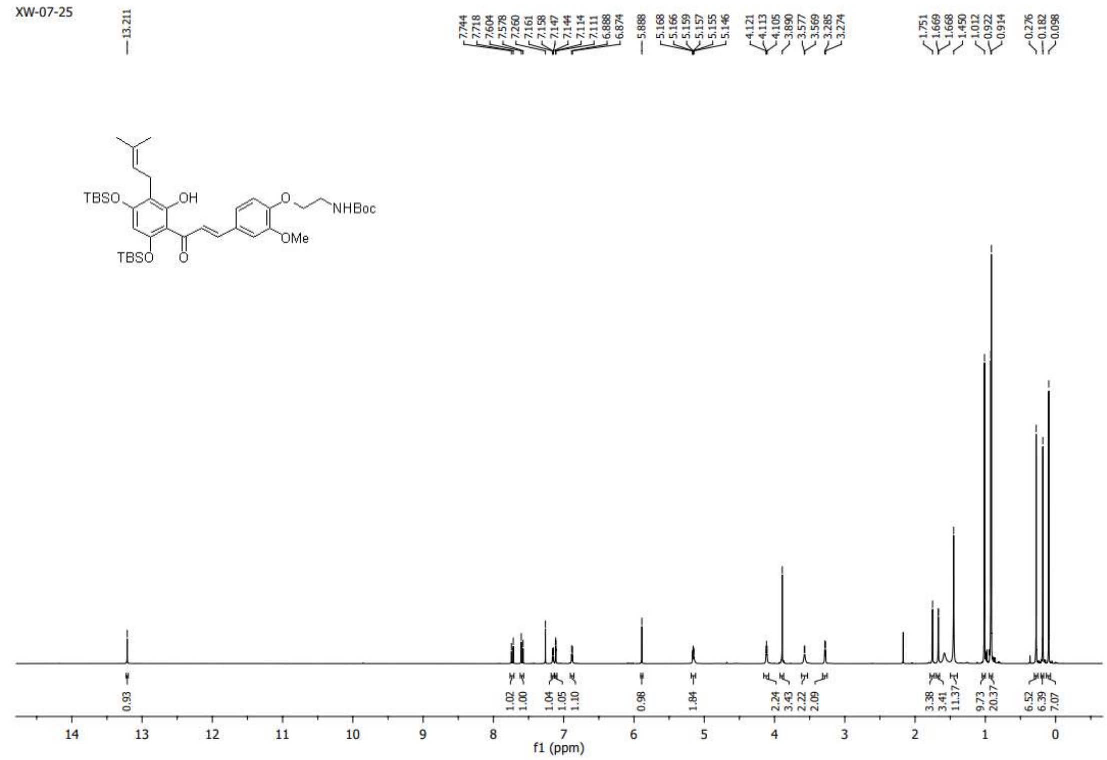

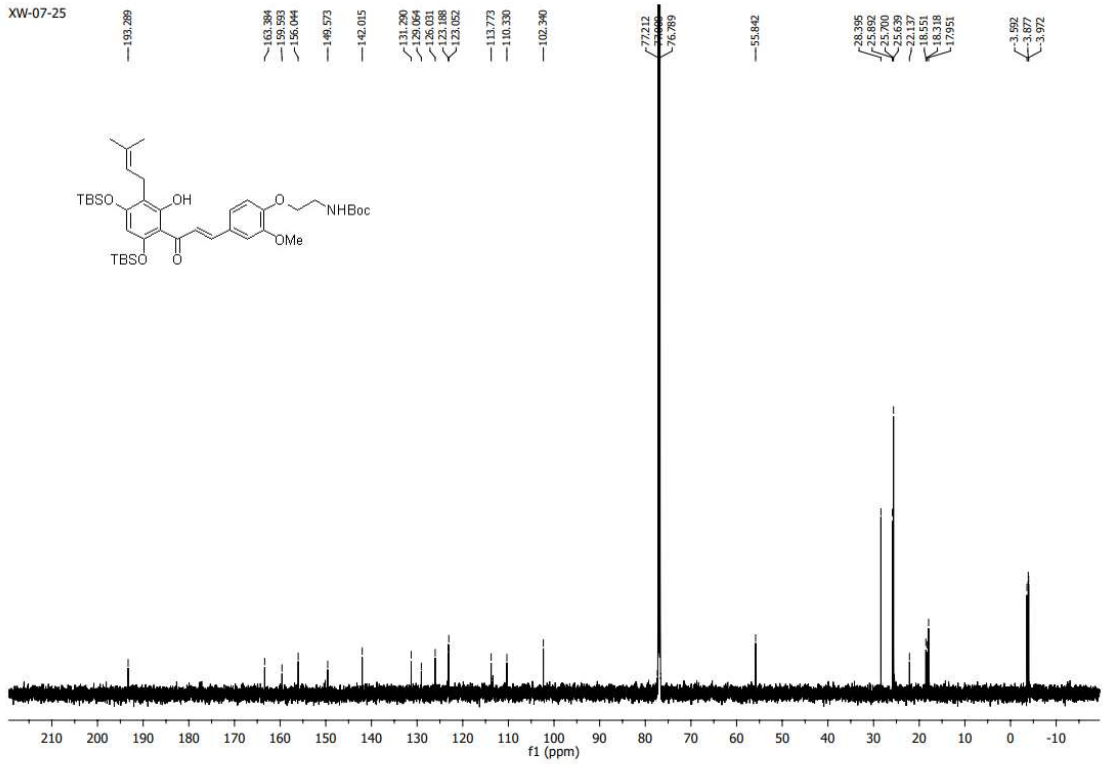

**Fig. S18**. $^{1}H$ and $^{13}C$ NMR Spectrum of Compound **7**

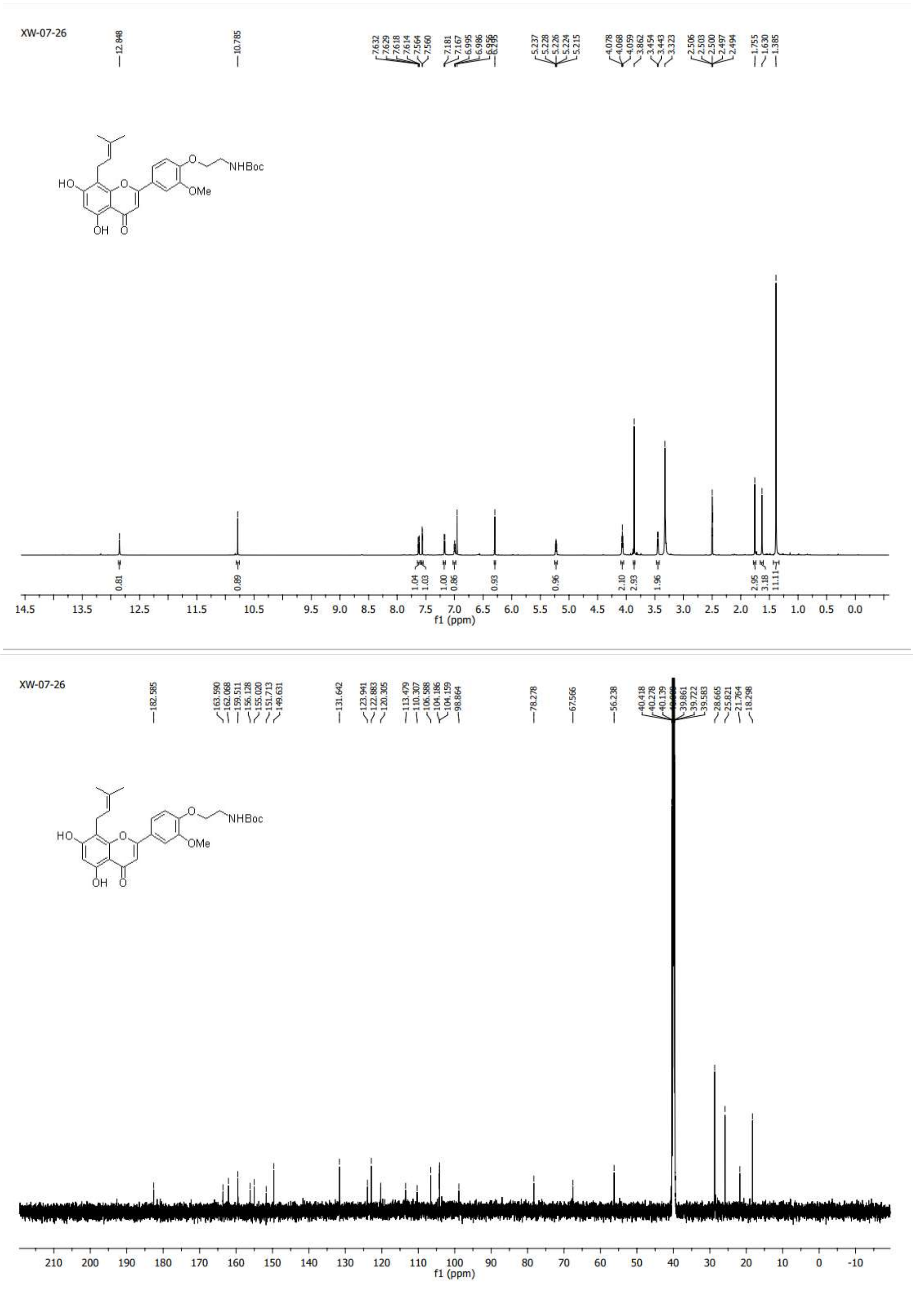

**Fig. S10**. $^{1}H$ and $^{13}C$ NMR Spectrum of Compound **8**

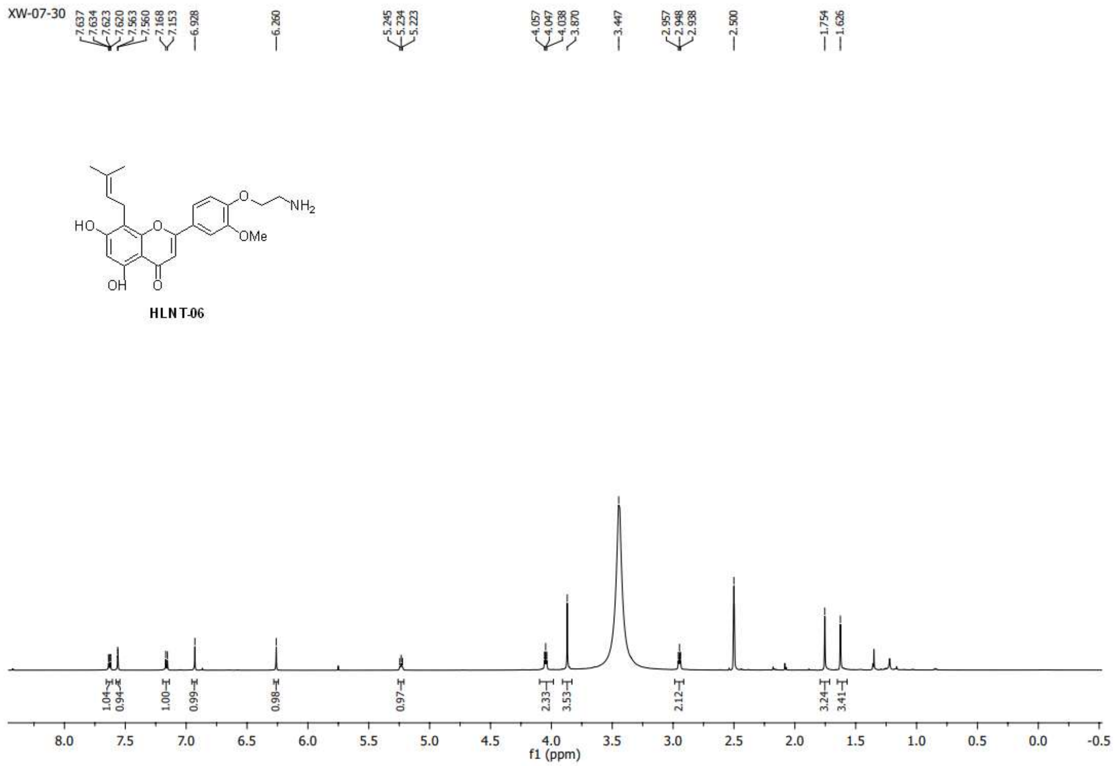

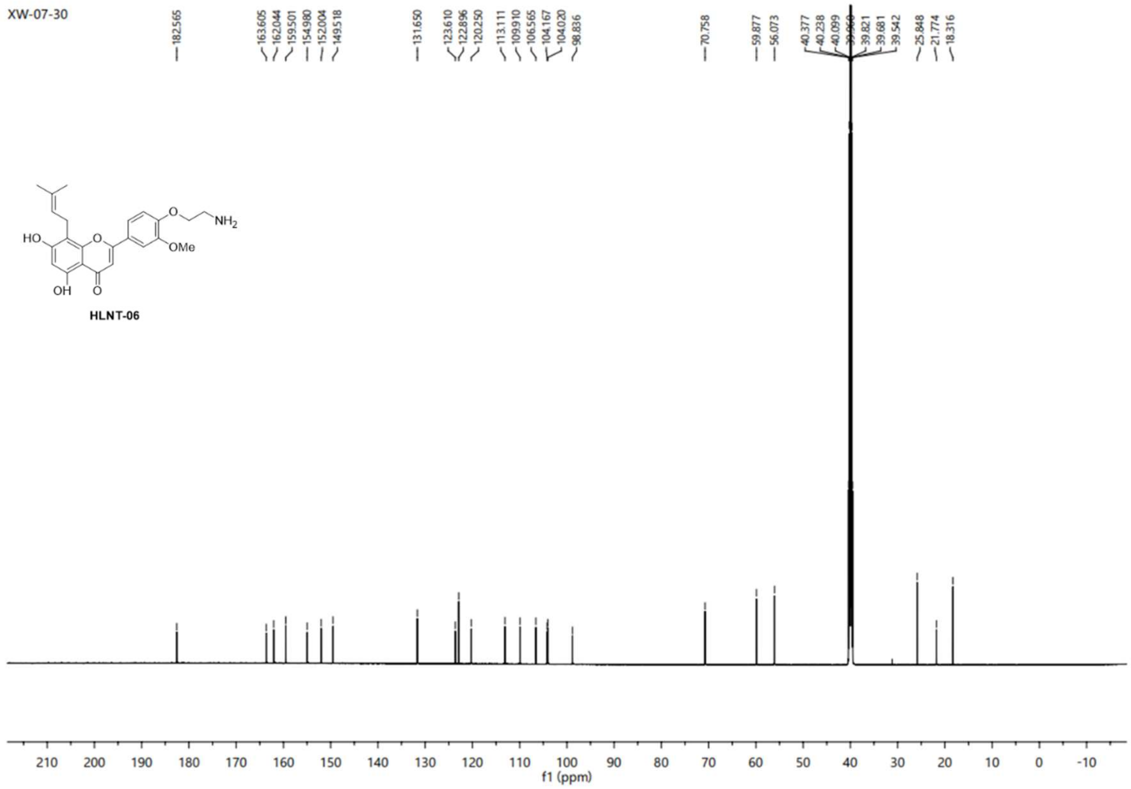

**Fig. S20**. $^{1}H$ and $^{13}C$ NMR Spectrum of Compound **HLNT-06**

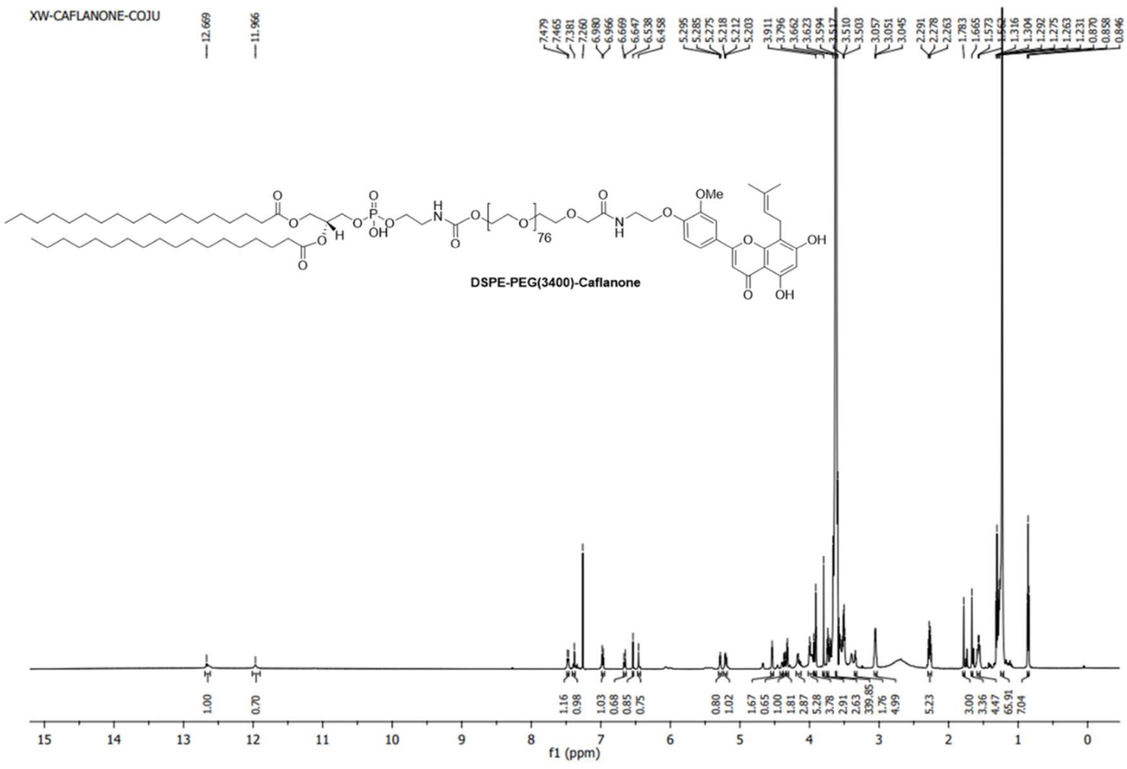


**Fig. S21**. [1]H Spectrum of **Caflanone-PEG(3400)-DSPE conjugate**

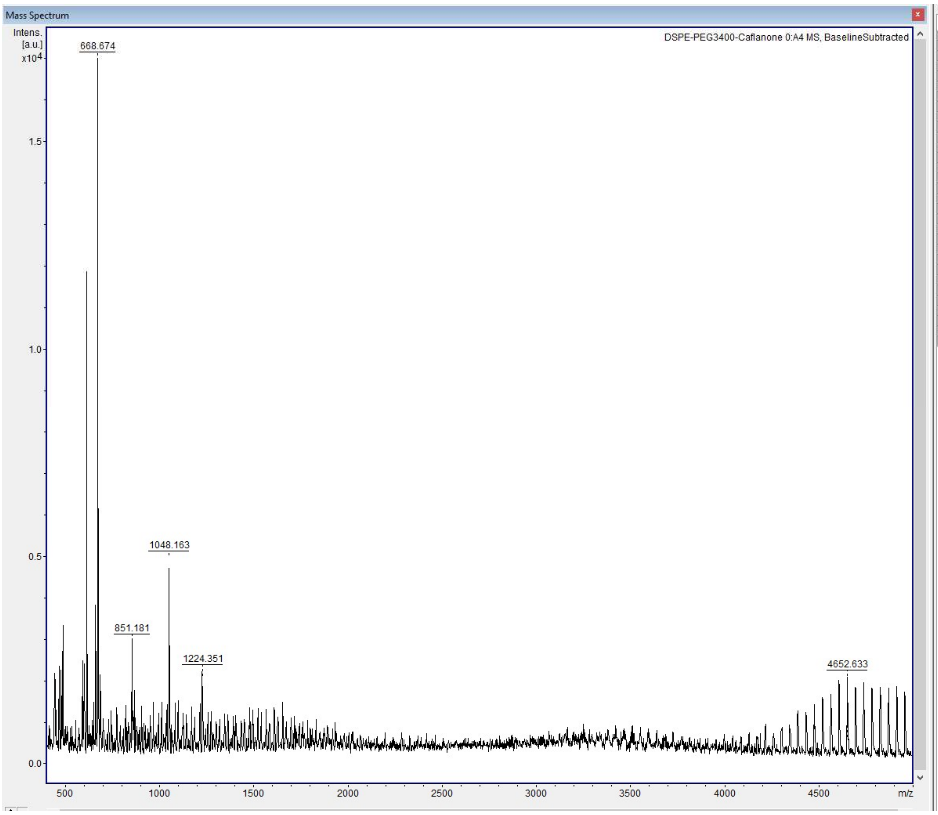


Mass List - 31 Peaks

| m/z | S/N | Quality Fac. | Res. |
|---|---|---|---|
| 438.182 | 9 | 1075 | 5700 |
| 441.056 | 9 | 590 | 8116 |
| 445.099 | 10 | 5715 | 9523 |
| 463.176 | 9 | 1082 | 742 |
| 474.313 | 10 | 737 | 8253 |
| 482.300 | 11 | 1620 | 2724 |
| 588.577 | 12 | 3712 | 12339 |
| 596.574 | 10 | 1336 | 9223 |
| 610.593 | 57 | 15747 | 12466 |
| 646.665 | 7 | 727 | 13424 |
| 656.492 | 18 | 14022 | 14908 |
| 658.682 | 7 | 503 | 11618 |
| 668.674 | 80 | 46798 | 13541 |
| 671.695 | 20 | 6007 | 13848 |
| 681.712 | 8 | 909 | 7389 |
| 851.181 | 14 | 11955 | 13286 |
| 857.952 | 6 | 287 | 18247 |
| 865.206 | 8 | 1667 | 15924 |
| 1048.163 | 23 | 6317 | 14483 |
| 1224.351 | 9 | 1850 | 9888 |
| 4521.220 | 7 | 114 | 13219 |
| 4565.026 | 8 | 655 | 12979 |
| 4608.834 | 8 | 922 | 13196 |
| 4652.633 | 9 | 1015 | 12977 |
| 4696.422 | 8 | 1027 | 12703 |
| 4740.190 | 9 | 358 | 12766 |
| 4783.956 | 9 | 1002 | 13030 |
| 4827.717 | 8 | 1153 | 13190 |
| 4871.490 | 8 | 1098 | 12006 |
| 4915.241 | 8 | 255 | 12675 |
| 4958.989 | 7 | 787 | 12379 |

Activate Windows
Go to Settings to activate Windows.

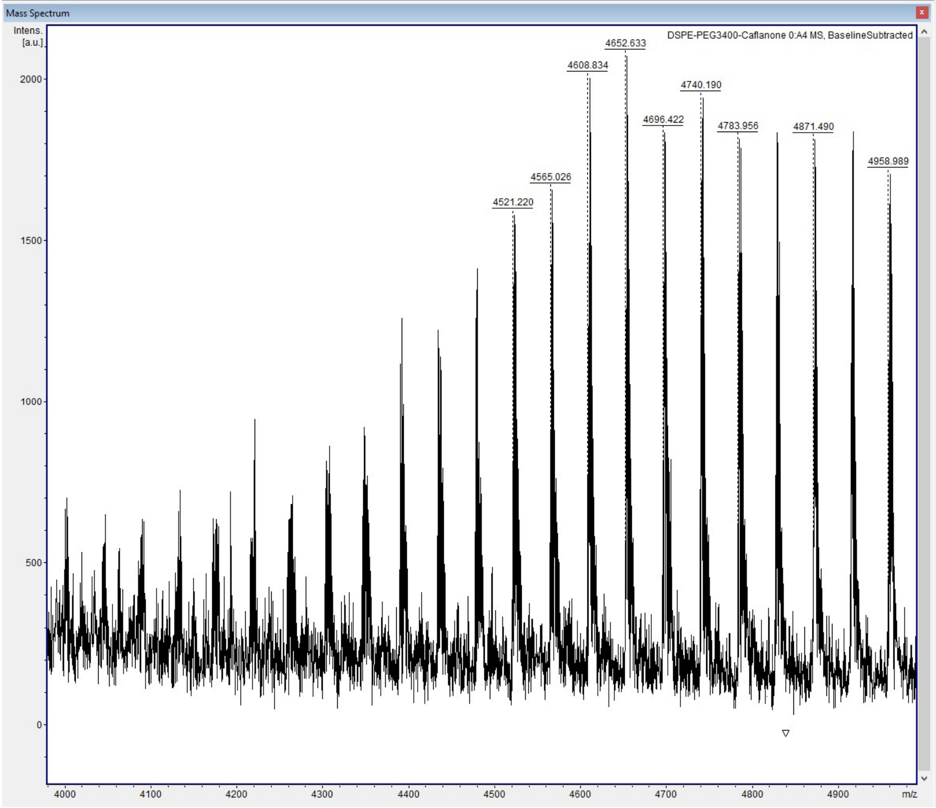


Mass List - 31 Peaks

| m/z | S/N | Quality Fac. | Res. |
|---|---|---|---|
| 438.182 | 9 | 1074 | 5700 |
| 441.056 | 9 | 590 | 8116 |
| 445.099 | 10 | 5716 | 9523 |
| 463.176 | 9 | 1082 | 742 |
| 474.313 | 10 | 737 | 8253 |
| 482.300 | 11 | 1619 | 2724 |
| 588.577 | 12 | 3712 | 12339 |
| 596.574 | 10 | 1336 | 9223 |
| 610.593 | 57 | 15745 | 12466 |
| 646.665 | 7 | 725 | 13504 |
| 656.492 | 18 | 14028 | 14909 |
| 658.682 | 7 | 503 | 11620 |
| 668.674 | 80 | 46798 | 13541 |
| 671.695 | 20 | 6009 | 13849 |
| 681.712 | 8 | 905 | 7368 |
| 851.181 | 14 | 11958 | 13295 |
| 857.952 | 6 | 287 | 18247 |
| 865.206 | 8 | 1666 | 15924 |
| 1048.163 | 23 | 6317 | 14483 |
| 1224.351 | 9 | 1850 | 9888 |
| 4521.220 | 7 | 114 | 13219 |
| 4565.026 | 8 | 655 | 12979 |
| 4608.834 | 8 | 922 | 13196 |
| 4652.633 | 9 | 1015 | 12977 |
| 4696.422 | 8 | 1027 | 12703 |
| 4740.190 | 9 | 357 | 12766 |
| 4783.956 | 9 | 1002 | 13027 |
| 4827.717 | 8 | 1152 | 13187 |
| 4871.490 | 8 | 1098 | 12006 |
| 4915.241 | 8 | 255 | 12675 |
| 4958.989 | 7 | 787 | 12379 |

Activate Windows
Go to Settings to activate Windows.

**Fig. S22.** MALDI Spectrum of **Caflanone-PEG(3400)-DSPE conjugate**
